# Loss of Fanconi anemia proteins causes a reliance on lysosomal exocytosis

**DOI:** 10.1101/2025.01.23.634631

**Authors:** Becky Xu Hua Fu, Albert Xu, Hua Li, Daniel E. Johnson, Jennifer R. Grandis, Luke A. Gilbert

## Abstract

Mutations in the FA pathway lead to a rare genetic disease that increases risk of bone marrow failure, acute myeloid leukemia, and solid tumors. FA patients have a 500 to 800-fold increase in head and neck squamous cell carcinoma compared to the general population and the treatment for these malignancies are ineffective and limited due to the deficiency in DNA damage repair. Using unbiased CRISPR-interference screening, we found the loss of FA function renders cells dependent on key exocytosis genes such as SNAP23. Further investigation revealed that loss of FA pathway function induced deficiencies in lysosomal health, dysregulation of autophagy and increased lysosomal exocytosis. The compromised cellular state caused by the loss of FA genes is accompanied with decreased lysosome abundance and increased lysosomal membrane permeabilization in cells. We found these signatures *in vitro* across multiple cell types and cell lines and in clinically relevant FA patient cancers. Our findings are the first to connect the FA pathway to lysosomal exocytosis and thus expands our understanding of FA as a disease and of induced dependencies in FA mutant cancers.

## Introduction

Fanconi anemia (FA) is a rare genetic disease that occurs in 1 in 300,000 live births. FA is caused by either inheriting biallelic germline loss-of-function mutations in one of the 22 genes (FANCA/C/D1/D2/E/F/G/I/J/L/M/N/O/P/Q/S/T/U/V/W) in the FA pathway or a monoallelic loss-of-function of RAD51 ^1^. The 23 genes associated with FA are all involved in a DNA repair pathway that detects covalently linked DNA strands (inter-strand crosslinks: ICL) and mediates repair through homologous recombination. Due to the deficiency in DNA repair machinery, FA patients have an increased risk in bone marrow failure (BMF)/ acute myeloid leukemia (AML). Although modern bone marrow transplant technology has allowed FA patients to overcome BMF and AML, FA patients have increased rates of solid tumors at remarkably young ages. Head and neck squamous cell carcinoma (HNSCC) are the most common solid tumor malignancy in FA patients with a frequency 500-fold to 800-fold higher in FA compared to the general population ^2^.

In addition, the inherent defect in DNA repair machinery in FA patients complicates treatment of FA patients’ cancers. The treatments include surgical removal and/or controlled low doses of chemotherapy, cross linking agents (cisplatin), and radiation. FA cancer patients usually suffer from severe radiation and chemotherapy toxicities even with low doses. Due to the limited options of treatment, the average life expectancy of FA patients is 20-30 years ^3^. The dire reality of the health issues for people with FA highlights the importance of having a more holistic understanding of the biology of FA pathway and the implications of its absence.

The majority of studies that characterize FA pathway function have focused on the critical role that FA proteins play in the repair of ICLs and in genome maintenance. However, noncanonical roles, including stabilization of replication forks and regulation of cytokinesis, have also been established ^4^. Additionally, functional genomics technologies have allowed for knowledge on even well studied pathways such as FA to be expanded. For example, unbiased genome-scale RNAi screens designed to identify genes that regulate autophagy unexpectedly identified FA genes and through further characterization it was shown that the FA pathway has novel functions in mitophagy^5,6^.

To further identify and characterize noncanonical functions and biology of the FA pathway, we performed genome-scale CRISPR-interference (CRISPRi) screens in isogenic FA pathway mutant (FANCD2) and wild-type (WT) cellular models to identify genetic dependencies induced by loss of the FA pathway activity. We unexpectedly found that genes related to lysosomal exocytosis are required for cell viability upon loss of FANCD2. Characterization of top hits such as SNAP23 and STX4, which are important for lysosomal exocytosis, led us to discover inherent lysosomal deficiencies in FA mutant cells. Our findings reveal an unexpected phenotype with the loss of FA pathway that advances our understanding of FA beyond a disease of DNA repair to include defects in quality control systems in lysosomal pathways that cause dysregulation of the major autophagy and lysosomal transcriptional regulator, which can render cells dependent on lysosomal exocytosis.

## Results

### Genome-scale CRISPRi screens identify lysosomal related genes as specific FA-deficient genetic dependencies

To characterize the genetic dependencies of an FA-deficient versus WT genetic background, we used CRISPR-Cas9 to engineer a FA deficiency (FANCD2-null) in a well-characterized head and neck cancer cell line (FaDu) (Figure S1). The FaDu cell line was chosen because the Fanconi Cancer Foundation (FCF) has available open source FANCA knockout (FANCA-null) and FANCA-null rescued with transgene (FANCA-null + Tg) isogenic FaDu and UM-SCC-01 (another head neck cell line) cell lines that complement our study design and provide convenient independent validation. We further engineered the parental FaDu and FANCD2-null cell models to stably express CRISPRi (dCas9-Krab) protein. We performed genome-scale CRISPRi screens targeting 18,905 protein coding genes in both FA mutant and WT FaDu CRISPRi cell lines and compared quantile normalized gene depletion scores between mutant and WT screen data to identify genes that are selectively required for cellular growth or survival in the FANCD2-null background (Figure 1a, see Methods for details).

**Figure 1.**
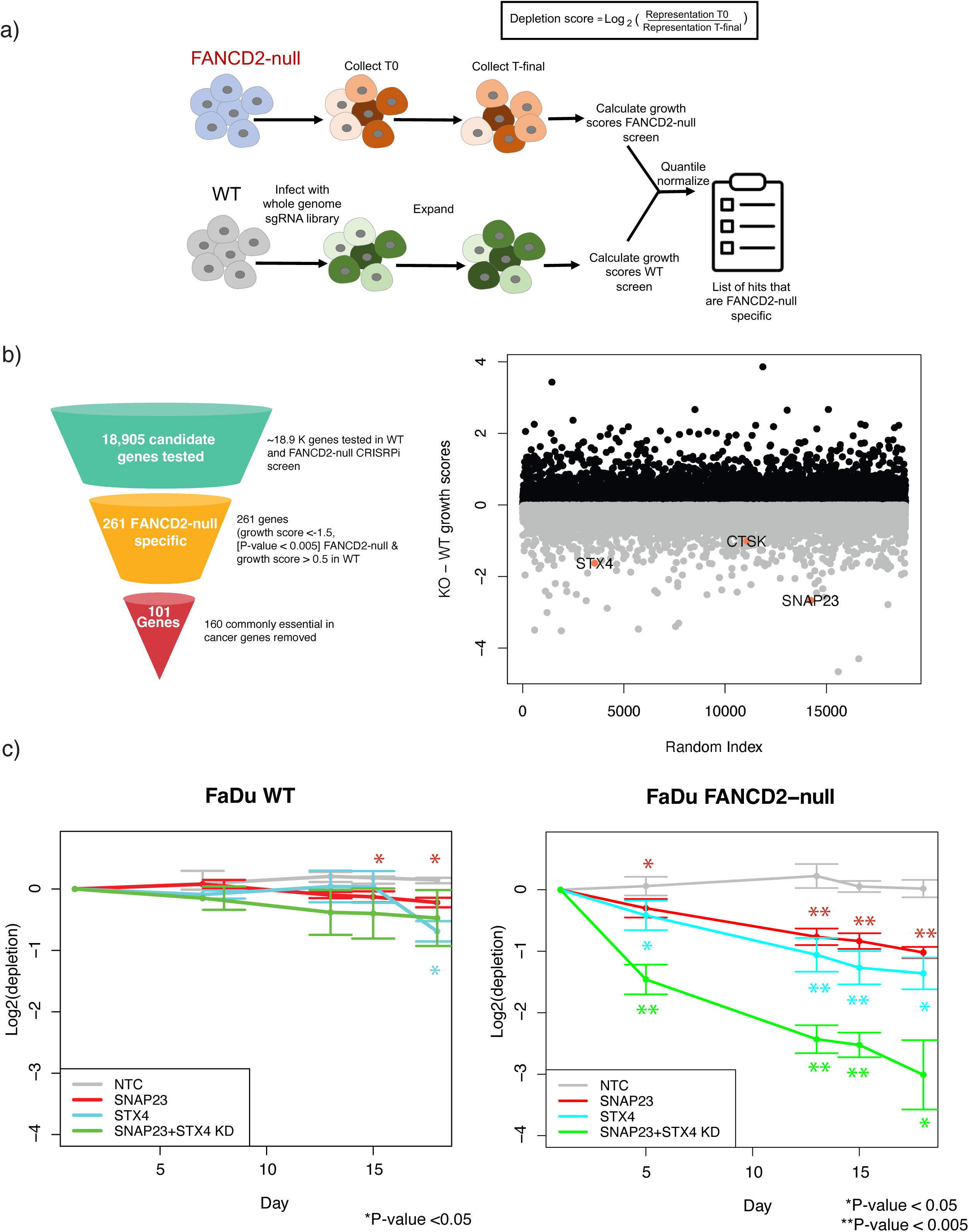
A CRISPRi screen nominates lysosomal biology as an induced dependency upon loss of FANCD2 a) A schematic of genome scale CRISPRi screens performed in FaDu WT and FANCD2-null cells. The depletion phenotypes from each screen were calculated, normalized, and compared between the WT and the mutant background to find candidate genes that were more depleted in the FANCD2-null background. b) The funnel diagram depicts the filtering to narrow down candidates from the screen (left). To the right is a scatter plot of CRISPRi screening results comparing FANCD2-null to WT conditions. Our screens nominated 101 candidate genes that are conditional dependencies in the FANCD2-null but not in the WT background after filtering for commonly essential genes. Genes that were involved in lysosomal and lysosomal exocytosis biology are highlighted in pink. Normalized depletion scores from the FANCD2-null background were subtracted from WT and plotted against a random index to visualize the spread of the differences between depletion scores. c) A graph showing a competitive growth-based assay analyzed over time to validate top FANCD2-conditionally essential CRISPRi screen hits in WT and FANCD2-null background FaDu cells. SNAP23 and STX4 were knocked down individually and together in both backgrounds and depletion was tracked over time (n = 3 as biological replicates; Mean ± STD, Unpaired two-tailed t-test was used to determine statistical significance).

We defined FANCD2-null conditionally essential candidate hit genes as having a depletion score lower than -1.5 and a p-value < 0.005 in the FANCD2-null screen and little to no growth effect in the WT background (depletion score>-0.5). The comparison between isogenic screens identified 261 genes that were conditionally essential in the FANCD2-null vs WT background (Table S1). Gene Ontology (GO) analysis using ShinyGO ^7^of the 261 unfiltered candidate genes specific to FANCD2-null background identified predictable pathways for a FA mutant, including cell cycle and chromosome segregation (Figure S2). This gene list was then filtered by removing all genes annotated as common essential by the Cancer Dependency Map (DepMap)^8^ (see Table S2 for common essential gene list that was used). The final FANCD2 conditionally essential candidate gene list consisted of 101 candidate genes (Figure 1b-left, Table S3). GO analysis of the 101 candidate genes showed no significant enriched pathways. Upon examination of the molecular functions of FANCD2-specific candidates, we noted top candidate hit genes involved in lysosomal and lysosomal exocytosis biology. For example, the lysosomal and lysosomal exocytosis related genes Cathepsin K (CTSK), Synaptosome associated protein of 23 kDa (SNAP23) and Syntaxin 4 (STX4) are highlighted (Figure 1b-right). Given that FA genes have been recently shown to play a key role in mitophagy, we hypothesized that these lysosome related hit genes which are FANCD2-null conditionally essential could be of considerable interest and point to how FA-deficiency broadly effects cell biology beyond canonical DNA repair defects. We confirmed that SNAP23-knockdown (KD) does not cause increase in DNA damage in FaDu FANCD2-null cells by quantification of protein levels of γH2AX (Figure S3).

### Loss of FA causes a reliance on lysosomal exocytosis

Lysosomes are important degradative organelles that play a crucial role in cellular homeostasis. Lysosomes not only contribute to clearance and recycling of cellular components but there has also been evidence to show that lysosomes and other organelles of autophagic and endo-lysosomal systems can be released for clearance and cell-to-cell communication and can play a role in plasma membrane repair through a process called lysosomal exocytosis. The process of lysosomal exocytosis involves the formation of a trans-SNARE complex made up of vesicle associated membrane protein 7 (VAMP 7), SNAP23, and STX4. The trans-SNARE complex allows for the docking and the fusion of the membranes of the lysosomes and endo-lysosomal vesicles to the plasma membrane which allows for the release of its contents ^9^. Lysosomal exocytosis is one of the three known methods of cellular clearance (autophagosome/amphisome secretion, exosomes). Studies have shown that cells with lysosomal damage respond by using migratory autolysosomes to expel autolysosomes via exocytosis.

To confirm that loss of FA activity induces a cellular dependency on lysosomal exocytosis through SNAP23, we validated our CRISPRi screen results using a mixed competition growth assay (see Methods). We observed that SNAP23 repression in FANCD2-null genetic background shows a statistically significant increased depletion compared to WT with non-targeting control (NTC)-KD in FaDu cells, ∼-1 vs ∼0.14 depletion score, respectively (Figure 1c). In addition, knockdown of SNAP23 in a FANCA-null background versus WT background in Cal33 cells, another head neck cancer cell line, recapitulated comparable results (Figure S4). Because lysosomal exocytosis has previously been shown to require SNAP23 as well as syntaxin 4 (STX4) and VAMP7 ^9^, we also measured the phenotype of simultaneous knockdown of SNAP23 and STX4. Dual knockdown of SNAP23 and STX4 resulted in a significantly stronger negative growth phenotype relative to single gene knockdown of either SNAP23 or STX4 in the FANCD2-null background but not in the WT background in FaDu cells (Figure 1c). These experiments validated our screen results and indicated that deficiency in the FA pathway induces a dependency involving lysosomal exocytosis. SNAP23 and STX4 control various intracellular trafficking pathways that could affect ability. SNAP23 has been previously characterized to form docking neuronal SNARE complexes with SNAP25 that function in calcium triggered exocytosis ^10–13^. Although SNAP25 is not a significant candidate in the screen and has no effect alone in knockdown validation experiments, the repression of SNAP23 and SNAP25 showed synergistic depletion in early timepoints compared to the single knockdowns in the FANCD2-null background (Figure S5). Together our results suggest that loss of FANCD2 function may render cells reliant on lysosomal exocytosis for survival.

Although we cannot rule out viability effects of the loss of SNAP23 is not due to its role in general trafficking, an analysis of various trafficking and general autophagy genes (25/27 genes) the FANCD2-null CRISPRi screening results show no significant effects in depletion score. The 2 genes had statistically significant depletion score: ATG16L1(gene necessary for autophagy by regulating membrane trafficking^14^) had ∼-0.96 depletion score while Rab5A (protein that regulates the movement of vesicles by facilitating internalization and fusion to early endosomes^15^) had ∼0.63 depletion score (Figure S6). The lack of effect of most general trafficking genes in the FANCD2-null background on the viability suggests that the SNAP23 knockdown phenotype we observe is unrelated to general trafficking.

Our findings suggest that deficiencies in the FA pathway result not only in the canonical DNA repair phenotype but can result in the reliance on genes involved in lysosomal exocytosis to maintain cellular homeostasis in certain cell types. Lysosomal exocytosis involves the secretion of lysosomal content by the fusion of the lysosome with plasma membrane. The fusion of the exocytotic lysosome into the membrane leaves a LAMP-1 scar on the cell surface. We used LAMP-1 luminal antibody and flow cytometry to quantify lysosomal exocytosis in FaDu WT and FANCD2-null cells with and without NTC and SNAP23 knockdown (Figure 2a). FaDu FANCD2-null cells with NTC-KD showed ∼1.5-fold increase in lysosomal membrane scars compared to WT background while both WT and FANCD2-null background with SNAP23 repression showed a decrease in overall lysosomal exocytosis. The FaDu FANCA-null cell line showed ∼2-fold increase in surface LAMP-1 and the FANCA-null+TG did not restore the lysosomal exocytosis rates to WT. Furthermore, lysosomal exocytosis of the FANCA/D2-null and FANCA-null +Tg in FaDu cells can be activated with ML-SA1, a chemical previously characterized to induce lysosomal exocytosis^16–18^, which indicates we are studying the effects on lysosomal exocytosis (Figure S7).

**Figure 2.**
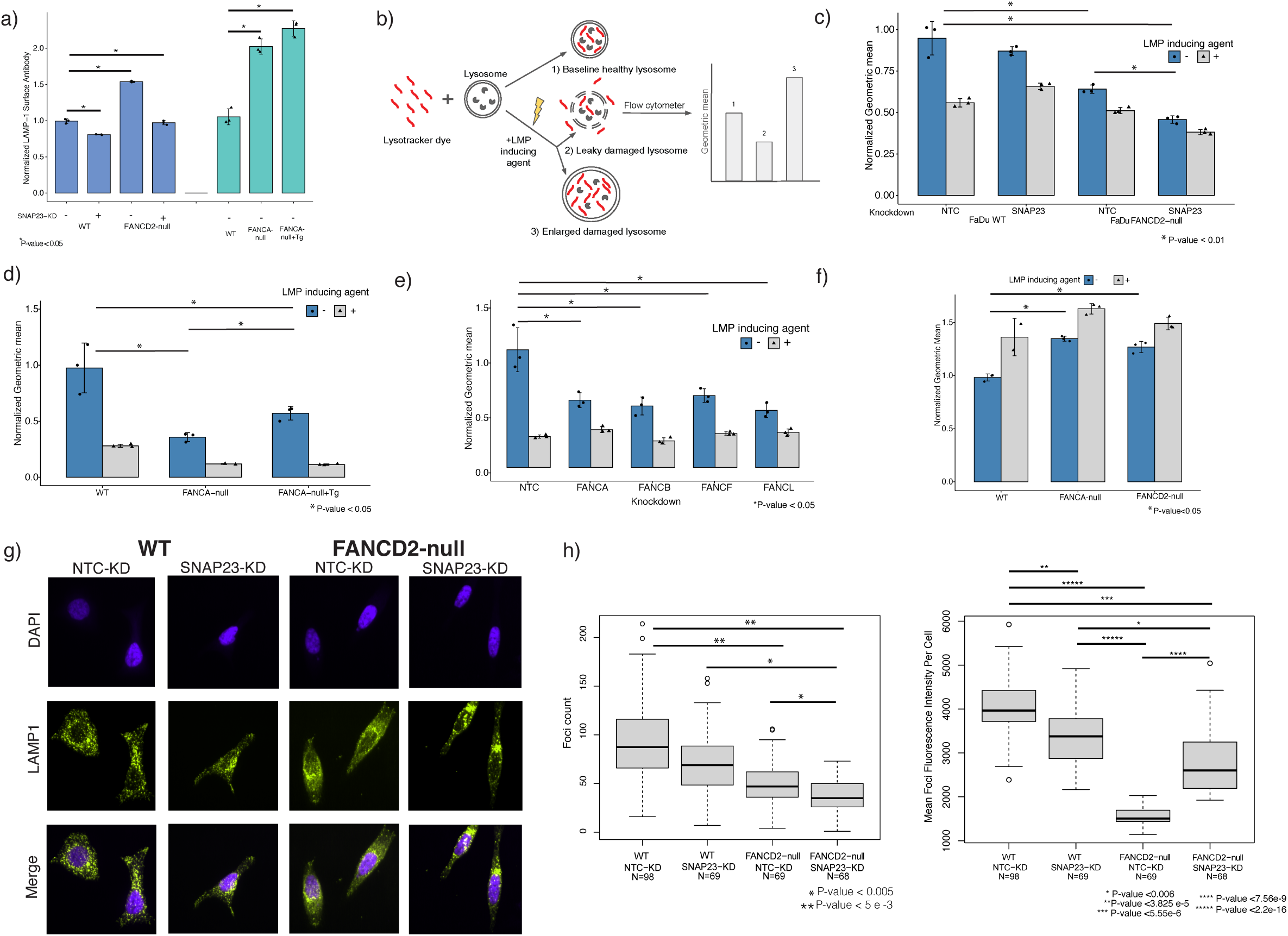
Loss of the FA pathway results in deficiencies in lysosomal health. a) Lysosomal exocytosis was quantified using antibody binding surface LAMP-1. Fadu WT and FANCD2-null background with NTC and SNAP23 repression and FA(A/A+Tg/D2) deficient knockout cell lines in FaDu. LAMP-1 scars on the plasma membrane were quantified by flow analysis (n = 3 as biological replicates; unpaired t-test was used to determine statistical significance). b) Graphical depiction of LysoTracker assay for assessing lysosomal membrane permeabilization. LysoTracker dye stains acidic compartments such as the lysosome. Depending on cell line, LMP damaged lysosomes can either be enlarged and have a higher fluorescent intensity or become damaged, leak, and/or dissolve which would be observed as lower fluorescence intensity using LysoTracker dyes. Using an LMP inducing agent (LLoMe or GPN) allows one to see the predicted dye retention of each cell line when a cell’s lysosomes are undergoing LMP. (All experiments in Figure 2 were performed in triplicate and reported as geometric mean ± STD, Unpaired two-tailed t-test was used to determine statistical significance.) c) A graph representing the lysosomal health as measured by LysoTracker dye of FaDu WT and FANCD2-null cells expressing control non-targeting control sgRNA (NTC-KD) and or SNAP23 (SNAP23-KD) in the presence or absence of LMP inducing agent (GPN). d) LysoTracker dye retention defects in FANCA-null and FANCA-null rescued with overexpression of a WT FANCA transgene. LLoMe was used was used as LMP inducing agent. e) The functional connections between FA pathway genes and lysosomal health were assayed in FaDu cells. FANCA/B/F/L knockdowns are nonessential in FaDu and were assayed for effect in lysosomal and endosomal health for the FaDu cell line. FANCA/B/F/L had a statistically significant effect on LysoTracker dye retention (decreases of ∼-2.44, - 2.33, - 2.86, -2-fold respectively). f) FITC-dextran can be used to monitor early changes in lysosomal pH during the LMP process in FANCA/D2-null FaDu cells. The fluorescence intensity of FITC-dextran is reduced in normal lysosomes (pH 4-5) and upon LMP, neutralization of lysosomes leads to increase in fluorescence intensity. FITC-Dextran is overloaded into lysosomes and when chased with fresh media and fluorescence was measured using flow cytometry. LLoMe treatment was used as a positive control. g) Lysosomal marker, LAMP-1 (green), was used to visualize lysosome number and morphology in FaDu WT and FANCD2-null with and without NTC or SNAP23 knockdown. Representative images are shown in Figure 2g and quantification is shown in Figure 2h. h) (Left) A plot of the distribution of the number of LAMP-1 foci per cell. The FaDu WT NTC-KD showed statistically significant decreases in LAMP-1 foci count compared to WT SNAP23-KD, FANCD2-null NTC-KD, and FANCD2-null SNAP23-KD. The open circles represent outliers defined by R base boxplot function as values > 1.5x outside the interquartile range. (Two-sided Mann-Whitney test was used to assess statistical significance) (Right)A plot of the distribution of the average intensity of LAMP-1 foci. FANCD2-null NTC-KD had statistically significant decrease in mean LAMP-1 foci count compared to the SNAP23-KD, the mean intensity of the LAMP-1 foci is partially rescued and comparable to the intensity seen in the WT SNAP23-KD. (Two-sided Mann-Whitney test was used to assess statistical significance)

In addition, we quantified lysosomal exocytosis with and without the loss of FANCA and FANCA-null + WT FANCA transgene rescue (FANCA-null +Tg) in UM-SCC-01. We found the LAMP-1 cell surface signal in the cell line with the FANCA-null to be increased compared to WT ∼2.7 -fold in UM-SCC-01 cells and is partially rescued with overexpression of FANCA transgene in FANCA-null background (Figure S8). The no rescue to partial rescue in FaDu and UM-SCC-01 cell lines could be due to non-endogenous expression levels of the FANCA transgene in the rescued cell lines (Figure S9). Overall, cells that have lost the FA pathway show increased rates of lysosomal exocytosis.

### Loss of the FA pathway exhibits defects in lysosomal health that result in lysosomal membrane permeabilization

#### Loss of FA pathway exhibits defects in lysosomal health

To maintain homeostasis, lysosomes break down and recycle or discard cellular proteins and products, and lysosomal exocytosis is one pathway for the cell to release unwanted material from lysosomes to the outside of the cell. Previous studies show evidence in zebrafish models of 3 different lysosomal storage disorders (Mucolipidosis II [MLII], sialidosis, and Mucopolysarchridosis Type IVA [MPSIVA]) that enhanced lysosomal exocytosis can arise from different genetic causes of lysosomal dysfunction ^19^. We observed an increase in lysosomal exocytosis with the loss of the FA pathway and hypothesized lysosomal irregularities may be the cause for this observed phenotype. If lysosome dysfunction creates a reliance on the lysosomal exocytosis pathway to remove damaged lysosomes, the loss of the secretion pathway can cause lysosomes to become overwhelmed and break down and to release their contents into the cytosol through a process known as lysosomal membrane permeabilization (LMP)^20,21^ which can lead to cell death.

To functionally evaluate whether mutations in the FA pathway were altering lysosomal biology and potentially causing LMP, we used a previously established protocol to quantify lysosomal health and LMP using a lysometric dye, LysoTracker, that stains acidic compartments within a cell, such as lysosomes and endosomes^22^ (See Methods). LysoTracker dye can be used to quantify the endosomal/lysosomal state of normal WT cells to compare to cells in question. During LMP, protons leak through endosomal/lysosomal membranes which results in pH gradient changes that can be quantified using lysometric dyes such as LysoTracker. One potential result of LMP is lysosomes enlarge and will hold more lysometric dye and have a higher fluorescent intensity. Another result of LMP is lysosomes are damaged to the point of leaking protons and their inner contents which causes a pH gradient that can be captured as lower lysometric dye fluorescence intensity within the lysosome (Figure 2b) ^23–26^. Whether a cell line will have compromised lysosomes that swell or disintegrate is somewhat poorly understood and is cell line dependent, and thus an LMP inducing agent such as (GPN or LLoMe) is used as a positive control for such assays (Figure 2b)^22,23,27^. The damaged lysosomes that undergo LMP release damaging lysosomal proteins and enzymes in the cell that cause cellular damage and eventually lead to cell death.

We observed that KD of SNAP23 in WT FaDu cells has a non-statistically significant effect on retention of lysometric dye relative to the non-targeting control and the LMP inducing agent caused lower retention of the dye (Figure 2c). By contrast, FANCD2-null FaDu cells expressing a non-targeting control (NTC) sgRNA showed decreased lysometric dye retention in the lysosomes under baseline conditions relative to WT control cells suggesting an induction of LMP in the absence of FA-activity (Figure 2c). Repression of SNAP23 in FANCD2-null FaDu cells further decreased lysometric dye retention suggesting increased dependency on SNAP23 for maintenance of lysosome health in the absence of FA proteins’ function.

This lysosomal defect is also seen with the FaDu FANCA-null cells and could be partially rescued in by transgene overexpression (Figure 2d) and fully rescued in UM-SCC-01 cells (Figure S10). As mentioned before, the partial rescue observed in FaDu cells could be due to non-endogenous expression levels of the FANCA transgene (Figure S9). Likewise, supplementing with overexpression of SNAP23 did not rescue the phenotype in the FaDu FANCA/D2 deficient cell line for LysoTracker dye retention and showed no difference in FANCA-null +Tg background (Figure S11).

We also tested the impact of loss of FA activity in different cell lines (FaDu, Cal33, UM-SCC-01, LN18, and LN229) on lysosomal health with and without the LMP inducing agent control (Figure S12). We observe a clear demarcation of the effect of LMP are cell line specific with LLoMe with the LysoTracker experiments: FaDu, Cal33, and LN18 with LMP inducing agent produce a phenotype of leaking/broken lysosomes while UM-SCC-01 and LN229 show a swelling of lysosomes. FaDu, Cal33, UM-SCC-01 are HPV negative head and neck cancer cell lines, while LN18 and LN229 are glioblastoma cell lines. Overall, the loss of FANCA in FaDu, Cal33, UM-SCC-01 and the repression of FANCD2 in LN18 had statistically significant defects in LysoTracker dye retention compared to WT conditions. Thus, we conclude knockout or knockdown of FANCA/D2 across diverse cellular models can induce lysosomal defect phenotypes.

#### The role of other FA proteins and DNA repair in lysosomal health

We then tested whether repression of other FA genes that are not essential for cell viability (FANCA/B/F/L) induced lysosomal and endosomal health phenotypes in FaDu cells. We found that repression of all FA genes tested had a statistically significant effect on lysosomal health relative to controls in FaDu cells (Figure 2e). We also tested whether FA gene function is important for lysosomal health in a non-cancer cell line. For these experiments we used RPE-1 cells, which are a TERT-immortalized retinal pigment epithelial cell line^28^ that is euploid and engineered to express CRISPRi. Here, we found that repression of FANCA and FANCD2 resulted in a significant elevated effect on the lysosomal dye retention suggesting lysosomal health also depends on FA gene function in non-cancer cells (Figure S13).

To test if other DNA repair genes similar phenotypes, we knocked down four additional DNA repair associated genes (XRCC4, FBXO42, ATMIN, and BRD8) that have minimal effects on growth in the FaDu WT CRISPRi screen (depletion score > -1). We found that the repression of XRCC4, FBXO42, ATMIN, and BRD8 also resulted in statistically significant decreases in the LysoTracker dye retention compared to the NTC-KD which suggests altered lysosomal states (Figure S14).

#### Lysosomal membrane permeabilization is observed with the loss of FA pathway

To further explore the lysosomal defects and LMP, we employed various other assays to characterize LMP and the general lysosomal health of the loss of function FA mutants. LysoSensor was used to quantify the pH of the lysosomes and a LMP inducing agent (LLoMe) was used as a positive control to lysosomes that are undergoing LMP. The pH of the lysosomes in FANCA/D2-null are, as expected of lysosomes undergoing LMP, more basic compared to WT in FaDu and UM-SCC-01 cells (Figure S15a). The FANCA-null+Tg does not rescue the acidity in the UM-SCC-01 cell line but does in the FaDu cell line (Figure S15a). The proteolytic/lysosomal activity measured using DQ-BSA Red and bafilomycin (an inhibitor of autophagosome-lysosome fusion) was used as a negative control. The proteolytic/lysosomal activity is higher in FANCA/D2-null mutants in FaDu and UM-SCC-01 cell lines and the activity is not rescued in the FaDu or UM-SCC-01 cell lines with the overexpression of a WT FANCA transgene in the FANCA-null background (Figure S15b).

To further characterize LMP in FA-mutant cells we used previously established protocols for pulse chase experiments with fluorescein isothiocynante-dextran (FITC-Dextran)^29,30^. Fluorescein isothiocyanate (FITC) conjugated to dextran can be used to monitor early changes in lysosomal pH during the LMP process. The fluorescence intensity of FITC-dextran is reduced in normal lysosomes (pH 4-5). Upon LMP, neutralization of lysosomes leads to increase in fluorescence intensity. FITC-Dextran is overloaded into lysosomes and when chased with fresh media. FITC conjugated to dextran can be used to monitor changes in lysosomal pH during the LMP process and LLoMe treatment was used as a positive control to simulate the results of cells in the assay that are undergoing LMP. Cells undergoing LMP will show increases in fluorescent intensity as expected with LMP agent which can be measured by either flow cytometry or imaging^31^. The FANCA/D2-null mutants showed statistically significant increase in FITC-Dextran fluorescent intensity in FaDu and UM-SCC-01 compared to WT cells (Figure 2f, Figure S16).

Another signature characteristic of cells undergoing LMP is the translocation of Galectin-3 (LGALS3) from the cytosol to the lysosomal lumen ^32–34^. Lysosome immunoprecipitation (Lyso-IP) was performed on the WT and mutant FaDu cell lines and the amount of LGALS3 was quantified using immunoblotting. An increase of LGALS3 was found in the lysosomes of the FANCA/D2-null mutants which corroborate the previous results (Figure S17). Another signature of damaged lysosomes undergoing LMP is the increase in K48-linked ubiquitination marks on lysosomes which signal engulfment by autophagosomes and recycling^35^. We found the lysosomes in the FaDu FANCA/D2-null cell lines had in increase in K48 ubiquitination (Figure S18). Overall, the data suggests that the defects in lysosomal health leads to LMP in cells deficient in the FA pathway.

From our data, we hypothesized that FA deficient cells may be more sensitive to chemical perturbations that cause lysosomal damage. There are well characterized chemicals that can cause lysosomal damage and LMP such as chloroquine (CQ). CQ preferentially accumulates in acidic organelles (lysosomes, endolysosomes, and Golgi), de-acidifies the luminal pH in acidic organelles, acidifies the cytosol and can cause LMP^36,37^. Consistent with this hypothesis, we observed that FA-mutant cells are more sensitive to CQ than isogenic control cells in clonogenic cell survival colony formation assays. Specifically, we found that FANCA and FANCD2-null had increased sensitivity to CQ compared to WT in FaDu cells, 2.6 and 1.78-fold increase respectively (Figure S19). Cal33 FANCA-null similarly was more sensitive to CQ than WT cells (∼1.75-fold sensitivity) and UM-SCC-01 FANCA-null had a ∼1.26-fold change in sensitivity relative to WT cells that was not statistically significant.

#### Loss of FA pathway causes differences in lysosomal morphology

To confirm the lysosomal defect, we used a lysosomal marker (LAMP1-lysosomal associated membrane protein 1) to visualize lysosomal morphology in WT and FANCD2-null FaDu cells with and without NTC or SNAP23 knockdown (Figure 2g-h; Figure S20 includes all data with LMP inducing treatment). Previous studies have shown that lysosomes and acidic organelles that undergo LMP lose puncta and staining fluorescence^38,39^. In this assay, we stained and imaged LAMP-1 and then used NIS-Element GA3 software to compile the images into a Z-stack to identify LAMP-1 foci for quantification of foci count, volume, and fluorescence intensity (see Methods for details). We observed a statistically significant decrease (∼-1.86-fold) in LAMP-1 foci count in FANCD2-null+NTC-KD compared to WT+NTC-KD (Figure 2h). The SNAP23-KD in FANCD2-null had the lowest foci count in all conditions (∼-2.5-fold decrease compared to WT NTC-KD) (Figure 2g-h). Impairing lysosomal exocytosis can cause lysosomes to accumulate damage and disintegrate which would lead to the statistically significant lower count of LAMP-1 foci seen following NTC and SNAP23 knockdown in the FANCD2-null background. We also observed the mean intensity of the LAMP-1 foci per cell is partially rescued in the FANCD2-null SNAP23 knockdown relative to the FANCD2-null NTC-KD condition (Figure 2g-h). The partial rescue of LAMP-1 mean foci intensity of FANCD2-null+SNAP23-KD is comparable to WT+SNAP23-KD levels. We hypothesize that the defective lysosomes in FANCD2-null NTC-KD cells have lower LAMP-1 count due to damaged lysosomes but retain the ability to secrete damaged lysosomes. However, the loss of SNAP23 on top of the loss of FANCD2 results in the impairment of cells’ ability to regulate lysosomal damage through release of damaged lysosomal contents, which can lead to lysosomal rupture and degradation^40^. Thus, the FANCD2-null SNAP23-KD results in decrease foci count and the semi-rescue of the LAMP-1 mean foci intensity due to the inability of the damaged lysosomes to be secreted.

### The loss of FA results in dysregulation of autophagy

FA proteins have a known role in mitophagy ^41–44^ and our findings of deficient lysosomes upon loss of FA proteins suggests a potential defect in general autophagy. Previous studies have shown that DNA damaging agents such as camptothecin ^45^, etoposide and temozolomide^46^, p-Anilioaniline^47,48^, and ionizing radiation^48^ arrest cell cycle and initiate autophagy. These findings show that even temporary activation of DNA damage can cause induction of autophagy. To explore potential autophagy deficiencies in the context of the FA pathway, we assessed mTORC1 activation by quantifying the levels phosphorylated 4E-BP1 (p-4E-BP1) and found an increase in p-4E-BP1 in FA mutants in FaDu cell lines (Figure S21).

The levels of p-4E-BP1 indicated dysregulation of autophagy, thus leading us to analyze the rate of autophagic degradation (autophagic flux)^49^. We assessed autophagic flux upon loss of FANCD2 and FANCA activity using a fluorescent protein probe (GFP-LC3-RFP-LC3ΔG). Upon expression, this probe is cleaved by endogenous ATG4 family proteases resulting in equimolar GFP-LC3 and RFP-LC3ΔG that cannot undergo lipidation ^50^. GFP-LC3 is degraded by autophagy and the RFP-LC3ΔG is released in the cytosol and serves as an internal control (Figure 3a). To measure autophagy, the ratio of GFP/RFP fluorescence intensity is measured with lower values indicating higher autophagic flux while higher values indicate lower autophagic flux ^50^.

**Figure 3.**
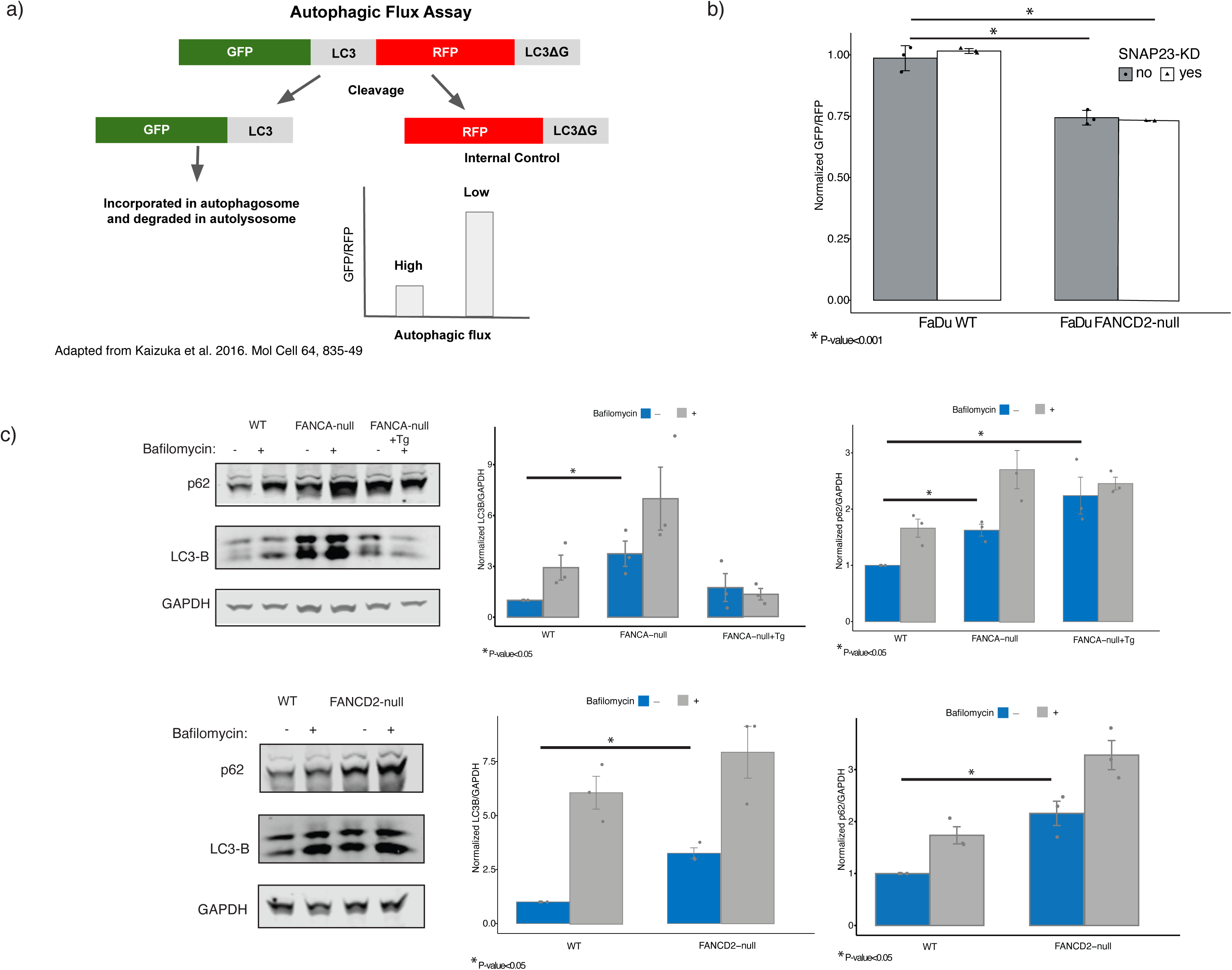
The loss of FA results in increased autophagic flux a) A schematic showing the GFP-LC3-RFP-LC3ΔG reporter assay used to assess the effect the loss of FANCD2 and FANCA on autophagic flux ^50^. The autophagic flux is interpreted by comparing the mean fluorescence intensity of GFP/RFP. A lower value indicates high autophagic flux while a higher value indicates lower autophagic flux. b) A graph quantifying autophagic flux in control cells and FANCD2-null cells with NTC or SNAP23 repression. Loss of FANCD2 with and without SNAP23-KD in FaDu show a ∼3.7-fold statistically significant increase in autophagic flux (P-value < 0.001) (n = 3 as biological replicates; Unpaired two-tailed t-test was used to determine statistical significance). c) Levels of LC3-B and p62 were measured using western blot for FANCA/FANCD2-null and FANCA-null+Tg. Biological replicates were done for each set of conditions and quantification are plotted on the right. (Unpaired two-tailed t-test was used to determine statistical significance)

Using this assay, we tested autophagic flux in FANCD2-null cells relative to control and upon SNAP23-KD in FANCD2-null or WT FaDu cells. We found that FANCD2-null cells with and without SNAP23 repression exhibited a statistically significant increase in autophagic flux (Figure 3b). Similarly, FANCA-null Cal33 cells also had increased autophagic flux compared to WT with and without SNAP23-KD (∼1.2-fold increase; p-value > 0.005), Figure S22). These results show that the loss of FANCA or FANCD2 leads to increased autophagic flux that is not additive or synergistic with the loss of SNAP23. This may suggest that SNAP23 is not directly involved in the FA-induced lysosomal defect.

To further characterize the autophagic flux changes in the FA mutants, we measured LC3-B and p62 levels. LC3-B is the insoluble, lipidated form of LC3 that is incorporated into the autophagosome while p62 is an autophagy receptor that binds to ubiquitin chains and signals targets to the autophagosome for degradation^49^. We found that the FA mutants have accumulation of LC3-B and p62 (Figure 3c). This suggests that the FA mutants exhibit increased flux through impaired degradation of cargo.

### Lysosomal signature is clinically relevant in FA cancers

To determine if our findings *in vitro* are observed in clinical data from patients with FA disease, we used publicly available transcriptomic patient data to identify genes that are differentially expressed between FA squamous cell carcinoma (SCC) patient tumors in comparison to sporadic head and neck squamous cell carcinoma HNSCC tumors ^51^. Ontology enrichment analysis on differentially expressed genes showed that genes that are upregulated in FA tumors predictably showed a statistically significant enrichment of DNA damage stimulus, DNA repair, DNA recombination, DNA double-strand break (Figure S23). The analysis of the down regulated genes showed a statistically significant enrichment of genes that are involved with the lysosome and endosomal processes: lysosomal membrane (p-value < 4.8 e-6), lysosomal lumen (p-value <8.0 e-5), endosome (p-value < 2.1 e-12), endosome membrane (p-value <8.2 e-8), lysosome (p-value < 3.0 e-9), and vesicle membrane (p-value < 1.7 e-7) (Figure 4a).

**Figure 4.**
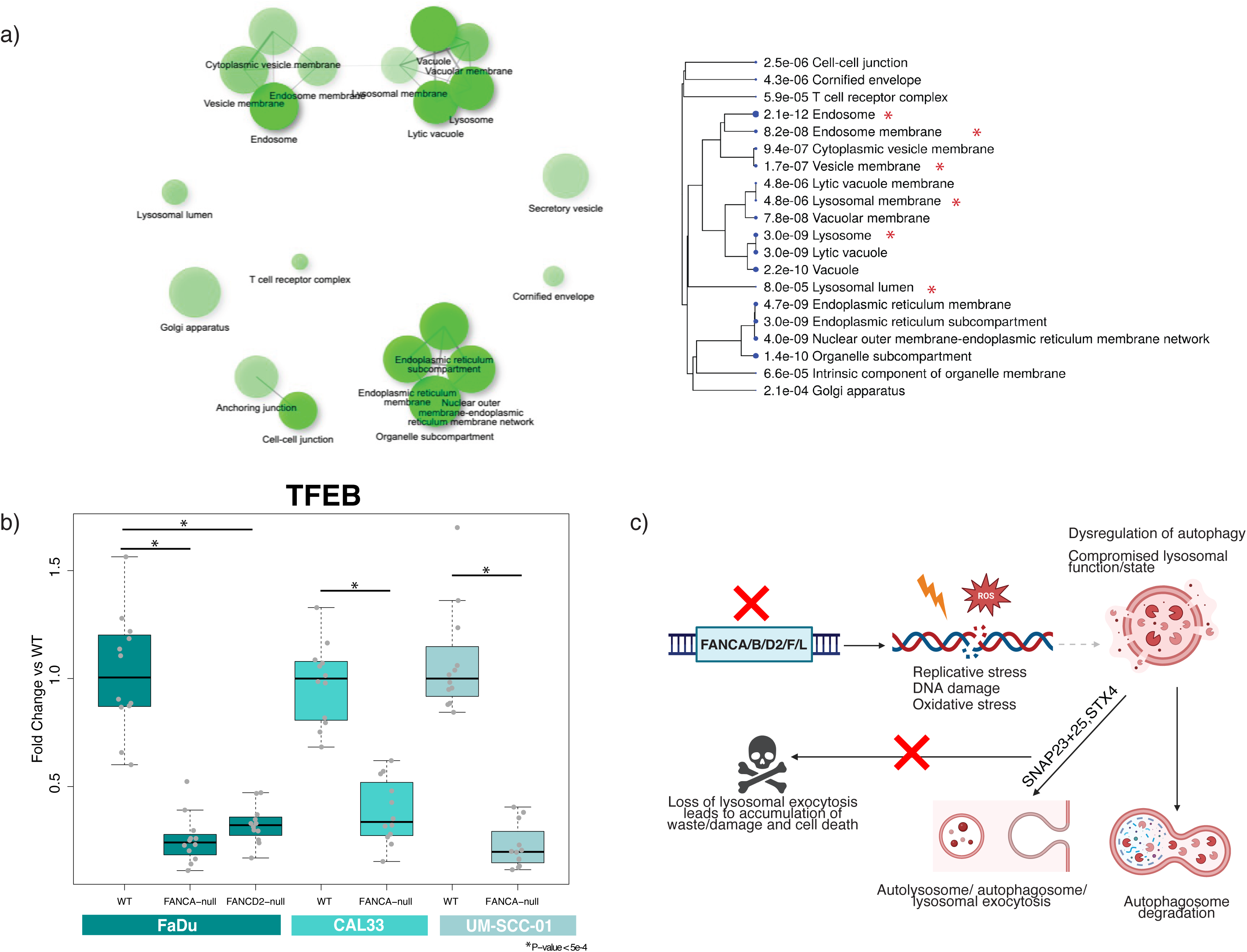
Transcriptomics of FA deficient patient samples demonstrate similar hallmarks of lysosomal and lysosomal exocytosis dysregulation. a) Publicly available patient transcriptomic data that analyzed differential expressed genes from FA SCC patient tumors in comparison to sporadic HNSCC (Webster et al. 2022) confirmed a lysosomal/lysosomal exocytosis phenotype. All genes that had a statistically differential gene expression difference (p-value < 0.005) by at least a 1.5-fold were used was input for GO analysis (ShinyGO v0.77)^7^. The results show enrichment in lysosome related functions that are highlighted with red asterisks. b) TFEB levels were measured using qPCR in FaDu, Cal33, and UM-SCC-01 FA-null mutants. c) A model of how deficiencies in FA pathway can lead to lysosomal defects and reliance on lysosomal exocytosis. The genomic stress that the loss of specific FA proteins causes replicative stress, DNA damage, and increase in oxidative stress. To combat these effects, there is dysregulation of autophagy and lysosomal biogenesis/health. This compromised state allows the cell to survive the loss of the FA pathway but renders the cell in a suboptimal state the is reliant on lysosomal exocytosis.

The FA SCC clinical data similarly showed a statistically significant decrease in expression in key lysosomal and lysosomal exocytosis related genes: TFEB and mucolipin 1 (MCOLN1) in FA SCC patient samples compared to sporadic HNSCCs (-1.71 log2 fold change (p-value < 2.01 e -9) and -2 log2 fold change (p-value < 8.46 e-19), respectively) ^51^. Notably, various lysosome specific proteins (LAMP-1/5) and enzymes (CTSB/D/F/G/S/W) were also found to be expressed significantly lower in FA SCC cancers (see Table S8 in Webster et al. 2022^51^). TFEB is transcription factor that specifically targets the Coordinated Lysosomal Expression and Regulation motif (CLEAR) to regulate lysosomal biogenesis and function through the transcriptional control of 96 lysosomal genes ^52^. By coordinating expression of lysosomal hydrolases, lysosomal membrane proteins, and autophagy proteins, TFEB can respond to lysosomal stress. In addition, TFEB is also known to play an important role in lysosomal exocytosis through Ca2+ signaling through MCOLN1. TFEB activates MCOLN1 to promote fusion of lysosomes to the plasma membrane to extrude lysosomal contents. TFEB is mainly localized in the cytoplasm and is translocated to the nucleus when activated ^52^. The cellular localization of TFEB is controlled by phosphorylation status by mTORC1. mTORC1 phosphorylates TFEB leading to cytoplasmic localization. Starvation and lysosomal stress that leads to mTOR inhibition leads to TFEB nuclear translocation and activation of the lysosomal and autophagy genes^53,54^.

We confirmed the signatures we found in the FA SCC clinical data in our cell lines by measuring mRNA expression of TFEB and MCOLN1 in multiple FA mutant and WT cell models. We observed substantial downregulation of TFEB (2.98-5-fold) across all FANCD2 and FANCA mutant cell models (FaDu, Cal33, UM-SCC-01) (Figure 4b). A statistically significant decrease of MCOLN1 was also found in FANCA-null in FaDu and UM-SCC-01 (Figure S24).

The localization of TFEB protein also plays a key role in the regulation of lysosomal biogenesis and autophagy. Phosphorylated TFEB is localized in the cytoplasm and upon dephosphorylation TFEB is translocated into the nucleus to induce transcription of the target regulatory genes. Thus, the ratio of cytoplasmic versus nuclear TFEB protein controls lysosome-autophagy pathway biology ^55^. Imaging of TFEB in FANCA-null FaDu and Cal33 cells showed the ratio of nuclear/cytoplasmic deviated in comparison to WT, showing a statistically significant increase in nuclear/cytoplasmic TFEB intensity the FaDu in FANCA/D2-null and a decrease in FANCA-null Cal33 cells (Figure S25a, c). In addition, immunofluorescence images confirmed that total TFEB protein levels quantified with TFEB fluorescence are lower in both FA deficient FaDu and Cal33 cell lines (Figure S25b, d). To determine if the lysosomal phenotypes were due to a reduction in TFEB levels, we transiently overexpressed TFEB and performed LysoTracker experiments. Consistent with the importance of proper TFEB localization, overexpression was not sufficient to rescue the lysosomal defect in FANCA/D2-null cells (Figure S25e). This suggests that loss of FA proteins impacts both levels and localization of TFEB with the latter being essential for proper lysosomal function.

While increased mTORC1 activity and decreased TFEB expression/nuclear localization are both associated with reductions in autophagy, we suspect that this could be due to the persistent stress and activation of mTORC1, which has been seen during persistent starvation^56^ leading to negative feedback on autophagy. TFEB and mTORC1 also have a complex relationship with TFEB as a target for mTORC1 and mTORC1 localization to lysosomes being impacted by TFEB-dependent processes such as endocytosis^57,58^. Overall, the FA patient gene expression data support our experimental findings that loss of FA pathway can cause defects in lysosomes and dysregulation of autophagy.

## Discussion

Patients with mutations in FA are at high risk of bone marrow failure/acute myeloid leukemia and have a 500 to 800 increase in HNSCC. The ability to treat these HNSCC malignancies is limited due to the inherent DNA repair deficiency in FA patients. FA proteins are primarily known to be involved in DNA repair and thought to cause diseases due to deficiencies in DNA repair and replicative stress which can also indirectly result in increased reactive oxygen species (ROS)^59,60^. The FA-null mutants for FaDu (FANCA/D2) and FANCA-null+Tg cell lines as expected, showed an increase in basal γH2AX which would be the catalyst of DDR that causes the changes in the autophagic state (Figure S3 & S26). In addition, we confirmed in our cell lines that FANCA and/or FANCD2-null cell lines (FaDu, Cal33, and UM-SCC-01) have increased ROS levels (Figure S27) and increased mitochondrial ROS (mtROS) (Figure S28).

Our findings reveal that deficiencies in the FA pathway can lead to significant lysosomal defects that may be a consequence of the inherent autophagic cellular state caused by DNA damage deficiencies and replicative stress. The crosstalk between DNA damage and autophagy has been reported to contribute to or prevent cell death depending on context^61,62^. Autophagy can mitigate DNA damage by regulating the recycling of the DNA repair proteins, mitochondrial quality control, controlling ROS production, ATP production, and cell death signaling^61^.

Our studies show that the cell lines lacking FANCA or FANCD2 have lysosomal defects and dysregulation in autophagy. The genotoxic stress and increase in ROS following loss of DNA repair activity may indirectly necessitate dysregulation of lysosomal state and autophagy to compensate and reach homeostasis to sustain viability. This compromised autophagic and lysosomal state may be dependent on controlling lysosomal biogenesis and autophagic state that relies on dealing with damage by using lysosomal exocytosis which results in dependency of SNAP23/STX4 for survival (see model: Figure 4c).

While we do not explore how defects in the FA pathway ultimately lead to these lysosomal phenotypes and dysregulated autophagy, it is unlikely that the connections are direct. Crosstalk between exogenous chemical and DNA damaging agents (camptothecin^45^, etoposide & temozolomide^46^, ionizing radiation^47^) have been shown to dysregulate autophagy. The autophagic changes that result from induced DNA damage response is temporary. In contrast, the loss of the FA pathway leaves cells less able to cope with endogenous forms of DNA damage (e.g. ICLs from aldehydes ^63^) and to protect replication forks^64–66^. These cells are likely in a somewhat persistent state of stress, and we speculate that the persistent loss of FA pathway in the cell alters the dynamics of autophagic processes and lysosomal quality control leading to a dependency on lysosomal exocytosis. Overall, our findings contribute to a broader understanding of the FA pathway and allows us to expand our understanding of how mutations in FA genes contribute to the pathology of the disease in the context of lysosomal health and lysosomal exocytosis.

## Methods

### Cell lines

Many of the cell lines used in this study were purchased from American Type Culture Collection (ATCC): FaDu, LN18, LN229, and RPE-1. FaDu were grown in RMPI 1640 (Gibco). LN18, LN229, and RPE-1 were grown in DMEM. All media was supplemented with 10% fetal bovine serum (FBS) with 1X penicillin-streptomycin-glutamine(pen-strep-glut) [Thermofisher #10378016] and grown in humidified 5% CO2 incubator at 37C. See Table S10 for list of cell lines.

FaDu FANCD2-null was made by electroporating Cas9 ribonucleoproteins (RNPs) into WT FaDu dCas9-Krab cell line using the Lonza Amaxa (ATCC HeLa presets). Candidate cell lines were isolated and established as clonal cell lines. Each candidate cell line was checked for the loss of function FANCD2 protein via immunoblotting.

FaDu and UM-SCC-01cell lines WT, FANCA-null, and FANCA-null+ WT FANCA transgene was generously donated by the FCF (www.fanconi.org; Fanconi Anemia Research Fund, n.d.). In addition, Cal33 WT and FANCA-null was also donated from FCF. UM-SCC-01 and Cal33 were grown in DMEM + 10% FBS + 1X pen-strep-glut.

### CRISPRi cell lines

CRISPRi cell lines were made according to previously published protocols^68,69^. Briefly, WT and FANCD2-null cell lines were infected with virus made from dCas9-KRAB construct (Addgene #217304) at low MOI (∼30%). Each cell line was sorted for dCas9-Krab using the BFP marker in the construct and validated for activity using an essential compared to NTC sgRNA in a depletion assay (see Validation/competition assay for details).

### Lentivirus production

All lentivirus made in this manuscript follows the Weissman lab protocol “Large scale lentivirus production protocol” (weissman.wi.mit.edu/crispr/). In summary the gag & pol (dR8.91), env (MD2G), lentiviral plasmid was transfected at an 8:1:8 ug ratio for 15 cm plate of 293T cells. The ratios were modified proportional to plate surface area for 10 cm plates.

### CRISPR screening and data analysis

Whole genome screens using the open source CRISPRi V2 library (Addgene 387 catalog #1000000091) were performed in FaDu WT and FANCD2-null background and published screening protocols were followed for CRISPRi screens ^69,70^. The genome scale library virus was generated and transduced at low MOI such that each cell expresses on average one sgRNA and thus one gene is knocked down per cell (∼30% infected). After 72 hours of infection, cells were selected on 5 ug/ml Puromycin. The sgRNA containing cells were expanded to establish 2-T0 and 2-T-final replicates each at 500x coverage. The 2 replicates were passaged for 14 days (∼10 population doublings) to allow for knockdown of genes to manifest as growth phenotypes in the presence or absence of FANCD2. Each replicate cell line was maintained at 500x coverage and 500x coverage was frozen down for the T-final. The samples were prepped using the Macherey Nagel NucleoSpin Blood kit and protocols. The concentration of each sgRNA in each sample (T0, T-final in WT and FANCD2-null) was quantified by sequencing and then sgRNA and gene level phenotypes were calculated for each screen.

The depletion scores were calculated using Weissman lab ScreenProcessing protocol ^69^. The collapsed gene depletion scores, raw counts and phenotype table can be found in for FANCD2-null can be found in Table S4-6 and for WT, Table S7-9. To compare depletion scores between WT and FANCD-null, depletion scores were quantile normalized using the R package preprocessCore ^71^. Quantile normalization was implemented to minimize technical variability. As stated above, FANCD2-null specific candidates of interest were filtered to have depletion score lower than -1.5 and p-value < 0.005 in the FANCD2 screen and no growth effect in the WT background.

### Validation/competition assays

To validate candidates, individual sgRNAs were cloned into the sgRNA vector for each gene of interest. The sgRNA vector used was either the BFP or mCherry sgRNA derived from pU6-sgRNA EF1Alpha-puro-T2A-BFP (Addgene #60955) ^69^. Each sgRNA candidate was cloned using BlpI and BstXI sites (detailed protocol found in https://weissman.wi.mit.edu/crispr/). WT and FANCD2-null cell lines were infected with the virus from each sgRNA candidate at a MOI ranging from 30-50%. The percentage of infected cells for each were tracked over 10-14 doublings using either a BFP or mCherry marker on the sgRNA plasmid using a flow cytometer (Attune NxT). The competition assay was performed using one sgRNA harboring the BFP and the other candidate with mCherry. WT and FANCD2-null cell lines were infected with each individual sgRNA and both. The percentages of each individual sgRNA and both were tracked via flow analysis. depletion scores were calculated by comparing log2 of the initial infection percentage and subsequent time points and end time point (see Figure 1a). Similarly, for validations the sgRNA candidate was tested against a NTC control, in this case a GAL4 sequence (5’-TTGGACGACTAGTTAGGCGTGTA-3’).

### Autophagic flux GFP-LC3-RFP-LC3ΔG assay

The GFP-LC3-RFP-LC3ΔG assay was established in the CRISPRi FaDu (WT & FANCD2-null) and Cal33 (WT & FANCA-null) using a modified pMRX-IP-GFP-LC3-RFP-LC3ΔG (Addgene #84572) ^50^. The GFP-LC3-RFP-LC3ΔG was cloned into a lentiviral vector (pFu-227). In short, lentivirus was made from the construct and infected at a low MOI in each cell line. Each infected cell line was then sorted for GFP and RFP and expanded. FaDu and Cal33 cell lines were then each infected with either NTC or SNAP23 sgRNA. After infection, GFP and RFP levels were measured via flow cytometer (Attune NxT).

### Lyso-IP experiments

Cell lines were made according to lysosome immunoprecipitation (Lyso-IP) protocol by Abu-Remaileh et al. 2017^72^. Cell lines were infected with pLJC5-Tmem192-3xHA (Addgene #102930) to introduce the lysosomal tag and was selected on Blasticidin (10 ug/ml). After cell lines are established, cell lines were expanded to 30 million cells for each cell line (FaDu-WT, FANCA-null, FANCD2-null) with and without 1 mM LLoMe treatment (30 minutes). Cells are rinsed twice with PBS and scraped with KPBS (136 mM KCl, 10 mM KH2PO4, pH 7.25 was adjusted with KOH). Cells were homogenized with a dounce homogenizer. The homogenate was centrifuged at 1000 x g for 2 min at 4C. The supernatant was removed and incubated with 50 ul of anti-HA magnetic beads (pre-washed with KPBS). The lysates were quantified using BCA assay (Thermofisher #23227) and 2000 ug lysate and HA beads were incubated at room temp on a rotator for 15 minutes. After incubation, the beads were washed 3 times with KPBS and resuspended in 50 ul of Laemmli buffer and used for western blotting (Galectin-3 [BD Biosciences 556904], LAMP-1[Cell Signaling D5C2P]).

### Western blotting

Protein from cells were lysed in RIPA buffer (Milipore 20188) with phosphatase and protease inhibitor (Thermofisher scientific #PI788445). Lysates were run on Bolt Bis-Tris Mini Protein gels (4-12%) on 0.45 um nitrocellulose membrane. Each experiment was performed at least twice, and representative images are shown (see Table S11 for list of antibodies).

### qPCR

At least 1 million cells were plated at ∼50-60% confluency overnight and harvested the next day. Cells were trypsinized and washed 2x with PBS. The RNA was extracted from each sample with the Macherey-Nagel RNA isolation kit (#740955.50). RNA was quantified and (∼100 ng) reverse transcribed using Thermo Scientific RevertAid cDNA synthesis kit (#K1621). For the reverse transcription 1 cycle of 25 C (10 min), 37 C (60 min), and 95 C (5 min) was followed. After reverse transcription, qPCR was done using Taqman Fast Advanced Master Mix (#4444963). The qPCR experiment and analysis were done using the Roche LightCycler 480-II. All primers were ordered from IDT Predesigned PrimeTime™ qPCR primers.

### LysoTracker/Lysosensor Flow Assays

Cells were counted using the Attune NxT and plated in a 12 well and allowed to adhere overnight. All FaDu derived lines were plated at 20K due to the cell lines apparent preference for higher density. While all cell lines derived from Cal33, LN18, LN229, UM-SCC-01, and RPE-1 were plated at 10K. Enough cells were plated for triplicates for each cell line plus and minus LMP induced treatment. For a positive control arms, original media is removed and replaced with 1 ml of 333 nM LLoMe (Cayman chemicals) or GPN (MedChemExpress) were added for 15-20 mins. GPN was used for early experiments but due to the short shelf life, powdered LLoMe reconstituted fresh was later used. After LMP inducing treatment, 50 nm LysoTracker Red (Thermofisher #L7528) or Lysosensor Green (Thermofisher #L7535) dye in each well for 30 minutes-1 hr. After incubation, wash cells 2x times with PBS and harvested using trypsin and resuspend in PBS+ 1% Fetal Bovine Serum. Samples were analyzed using the Attune NxT, using the YL2 channel for LysoTracker red and BL1 channel for Lysosensor. The geometric mean was calculated for all samples and a median was taken.

### Colony Formation Assays

Each cell line of interest was counted using the Attune NxT and 10K cells were seeded into each well of a 6 well. The cells were allowed to adhere overnight, and the next day varying doses of CQ resuspended in DMSO and a DMSO control was added. Fresh CQ was replace every 3^rd^ day for 3 times (9 days total). When assay in finished, cells were washed twice with PBS and 500ul of 0.5% crystal violet fixation solution (0.5 g crystal violet, 20ml of methanol, 80ml of water) was added in each well for 15-30 minutes. Colonies could be visualized after washing with water 3-5X times.

### DQ-BSA Pulse Chase Flow Assay

Cell lines were counted and plated at 20K in a 24-well plate for treated and untreated conditions and allowed to adhere overnight. For a negative control arms, original media is removed and replaced with 10 uM Bafilomycin (MCE #HY-1000558) and incubated for 2 hrs. After 2hr Bafilomycin treatment, master mix of DQ-BSA Red (Thermofisher #D120451) was incubated for 1 hr. After 1hr of DQ-BSA (1 ug/ml) chase, the media was changed and allowed to chase for 3 hr chase. When chase is finished, cells were washed with PBS 2x and then lifted and washed for flow analysis.

### Dextran-FITC Pulse Chase Flow Assay

Cell lines were counted and plated at 20K in a 24-well plate and allowed to adhere overnight. After overnight incubation, old media was removed and either new media with Dextran-FITC or Dextran-FITC + 1mM LLoMe (positive control) was added a 2 hr pulse. Dextran-FITC (AAT Bioquest #21700) 5000 ug/ml. After the 2 hr incubation the media was removed, and fresh media was added for an overnight incubation. The next day at 24 hrs, the cells were washed with PBS 2x and lifted and washed for flow analysis.

### ROS Assays

The Attune NxT was used to count cells to plate in 12 well plate. All FaDu derived lines were plated at 8K due to the cell lines apparent preference for higher density. While cell lines derived from Cal33 and UM-SCC-01 were plated at 5K. Enough cells were plated for cell line for triplicates plus and minus positive control. For the ROS assay CellROX Green (ThermoFisher #C10444) was used and for the MitoSOX Red (ThermoFisher #M36007) was used. Positive controls for CellROX were treated with hydrogen peroxide for 10 mins (1:1000 dilution of 30% peroxide in media). Positive controls for MitoSOX were treated with 50 uM mitoparaquat overnight. For the experiment, 500 nM CellROX or MitoSOX was incubated with the cell lines for 1 hour. After incubation, cells were wash with PBS 2x and lifted and resuspended with PBS+ 1% Fetal Bovine Serum cells for flow cytometry.

### Surface LAMP-1flow assay

About 100k cells were fixed with 4% paraformaldehyde for 15-30 minutes and washed twice with PBS. The cells are resuspended in PBS+10% fetal bovine serum (FBS) with no antibody (control) or with Santa Cruz Biotechnology anti-rat LAMP1-(H4A3 1:100; sc-20011) at 4 C for at least 60 minutes. All cells were then washed PBS and incubated with Alexa-647 conjugated anti-rat secondary for at least 30 mins at room temperature in PBS+10% FBS (1:1000) protected from light. Cells were then washed with PBS twice and resuspend in PBS+10% FBS for flow cytometry analysis (Attune NxT). The mean fluorescence of each condition was measured. To normalize for background fluorescence, each condition with the antibody+secondary was subtracted by the median of mean fluorescence of the corresponding condition’s control with only secondary. For plotting, all conditions were normalized by dividing by the median of the WT condition’s mean fluorescence. For ML-SA1 activation, cells were incubated with 50 mM ML-SA1 and processed as previously described.

### Immunofluorescence Microscopy and Analysis

#### Immunofluorescence Microscopy

Cells 150K cells for each cell line was plated on poly-L-Lysine coated cover slips and allowed to adhere overnight. For positive controls hydrogen peroxide (1:1000 dilution of 30% peroxide in media) was incubated with each slide for 10 minutes. All coverslips were fixed with 4% paraformaldehyde at room temperature (RT) for 15-30 minutes. Coverslips were then treated with blocking buffer (1.1% BSA, 0.1% Triton X-100 in PBS) for 15-30 minutes at RT. Each coverslip was incubated overnight in 4 C in 100 ul of 1:200 of LAMP-1 antibody (Cell Signaling Technology D4O1S anti-mouse). After primary antibody incubation, coverslips were washed 3x 5minutes with PBS + 0.1% Triton-100. Secondary antibody was added at a 1:500 dilution + DAPI for 1 hour at RT. Finally, coverslips are washed 2x 5 min with PBS + 0.1% Triton-100 and 2x5 min with PBS. Coverslips were sealed with Invitrogen ProLong Gold AntiFade and allowed to dry and were kept in –20 C and away from light. Images and data for this study was acquired at the Center for Advanced Light Microscopy-Nikon Imaging Center at UCSF on Spinning Disk confocal microscope.

#### Analysis

IF analysis was performed using GA3 in NIS-Element. Foci segmentation was done with NIS.ai in NIS-Element. Whole cell segmentation was done by applying ‘Grow Regions’ module in GA3 from nucleus area (segmented from maximum projected DAPI images by Otsu thresholding) to whole cell area. The count, volume, and intensity of LAMP-1 foci were assessed.

## Acknowledgements

We want to thank all members of the Gilbert and Grandis/Johnson lab for suggestions and discussions. We would like to thank Dr. Andrew Fire and Dr. Gavin Sherlock for helpful suggestions. We also want to thank SoYeon Kim at the UCSF’s Center for Advanced Light Microscopy for all guidance with imaging and analysis. We want to thank the Arc Institute Scientific Publications Team (Brian Plosky) for help and input. Brian Plosky gave invaluable insight and suggestions scientifically and during the writing process.

## Author contributions

B.X.H.F conceived and designed the study. B.X.H.F. and A.X. acquired data. B.X.H.F analyzed data. B.X.H.F and L.A.G. wrote the manuscript with input from all authors. L.A.G. acquired funding, provided equipment/materials, and advised the work. H.L. assisted with experiments supervised by D.J. and J.G.

