## Supplemental for "Loss of Fanconi anemia proteins causes a reliance on lysosomal exocytosis": Final_FA_Supplemental_2025.docx

Figure S1) FaDu FANCD2-null was validated using western blot. CRISPR-interference was validated in each line and clone 6 was chosen for the screen.

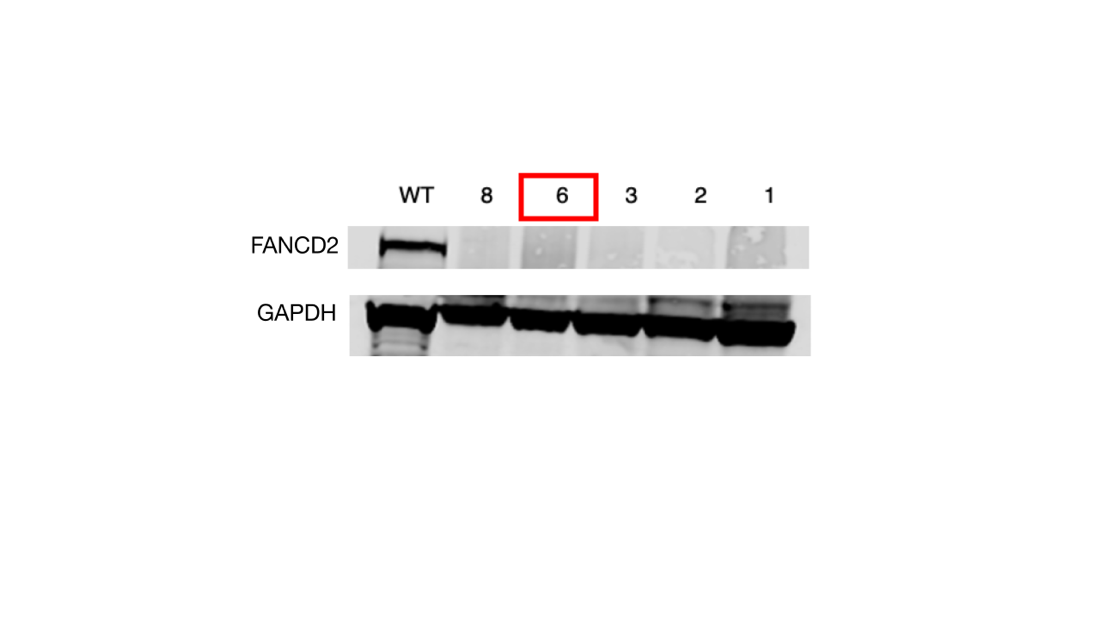

Figure S2) ShinyGO analysis for all 261 filtered at p-value<0.005 that are specific to the FANCD2-null background.

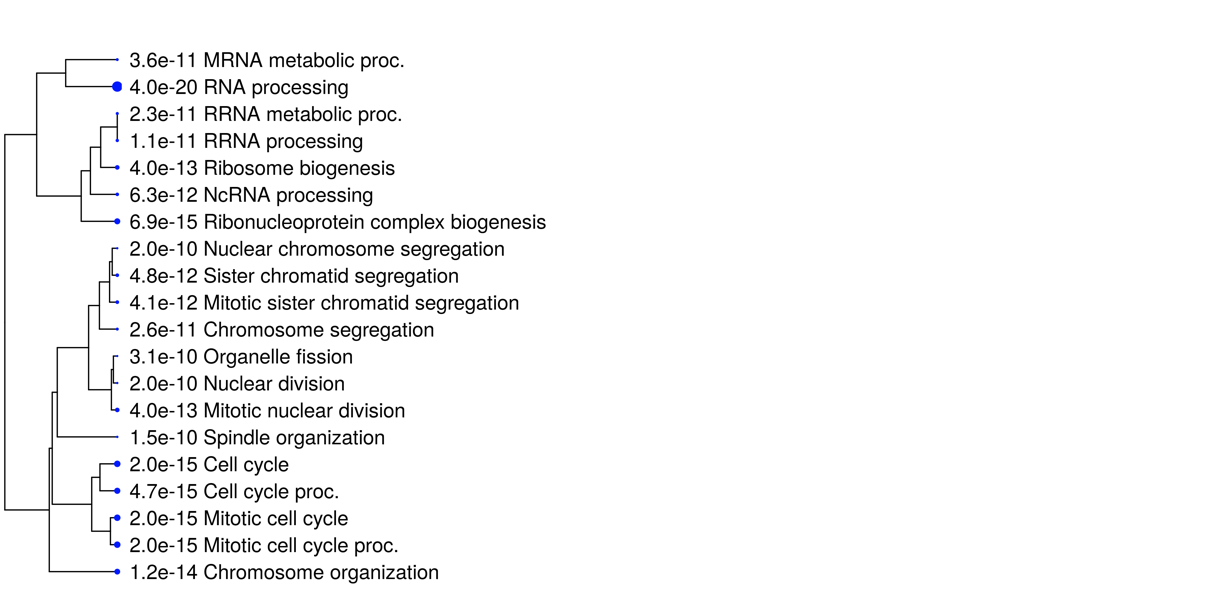

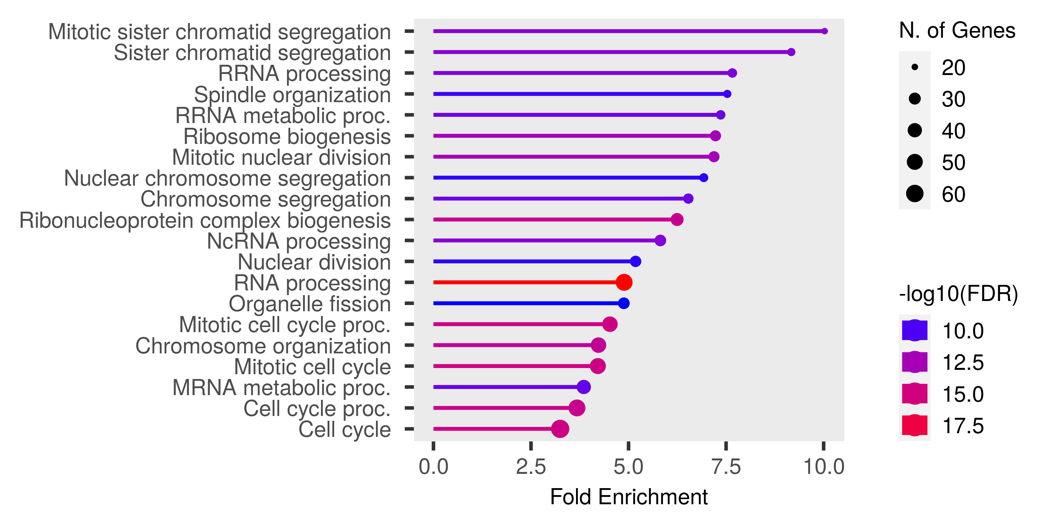

Figure S3) Protein levels of phospho-H2AX S139 (γH2AX, marker for DNA damage) were measured in FaDu WT and FANCD2-null with NTC and SNAP23 knockdown. Although the FANCD2-null has higher levels of γH2AX S139 compared to WT, the FANCD2-null with SNAP23-KD vs NTC-KD does not increase the levels of γH2AX.

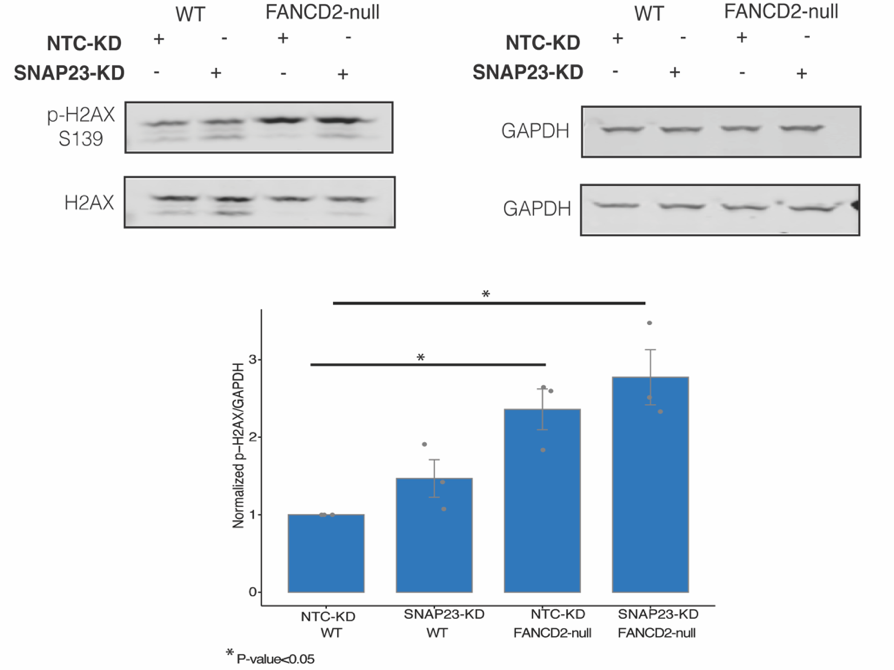

Figure S4) SNAP23-KD causes FANCA-null specific depletion in Cal33, a head neck cancer cell line. SNAP23-KD knockdown validated in Cal33 and Cal33 FANCA-KO CRISPRi cell lines. Knockdown of SNAP23 showed ~1.3-fold decrease in growth versus no effect in WT background (n = 3 as biological replicates; Median ± STD, Unpaired two-tailed t-test was used to determine statistical significance).

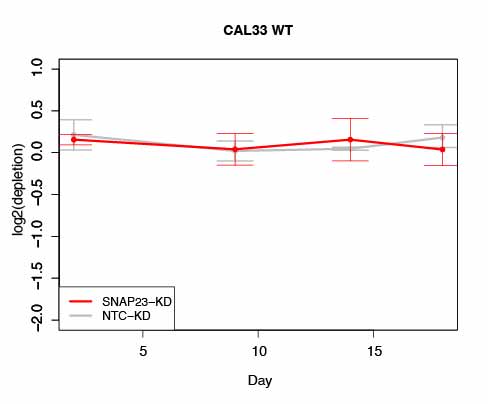

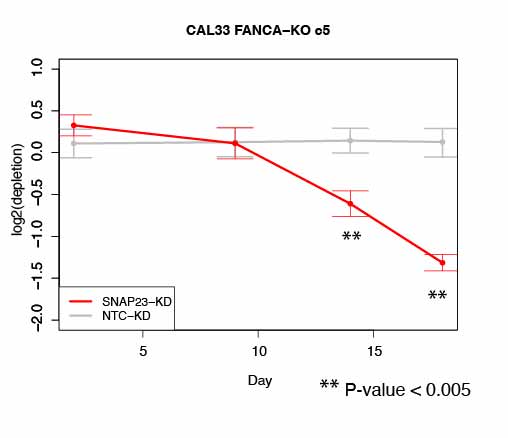

Figure S5) SNAP23 and SNAP25 repression shows a synergistic depletion in FANCD2-null background in the early timepoint. Repression of SNAP23+25 shows ~2-fold decrease in growth in FANCD2-null background compared to WT. (n = 3 as biological replicates; Median ± STD, Unpaired two-tailed t-test was used to determine statistical significance).

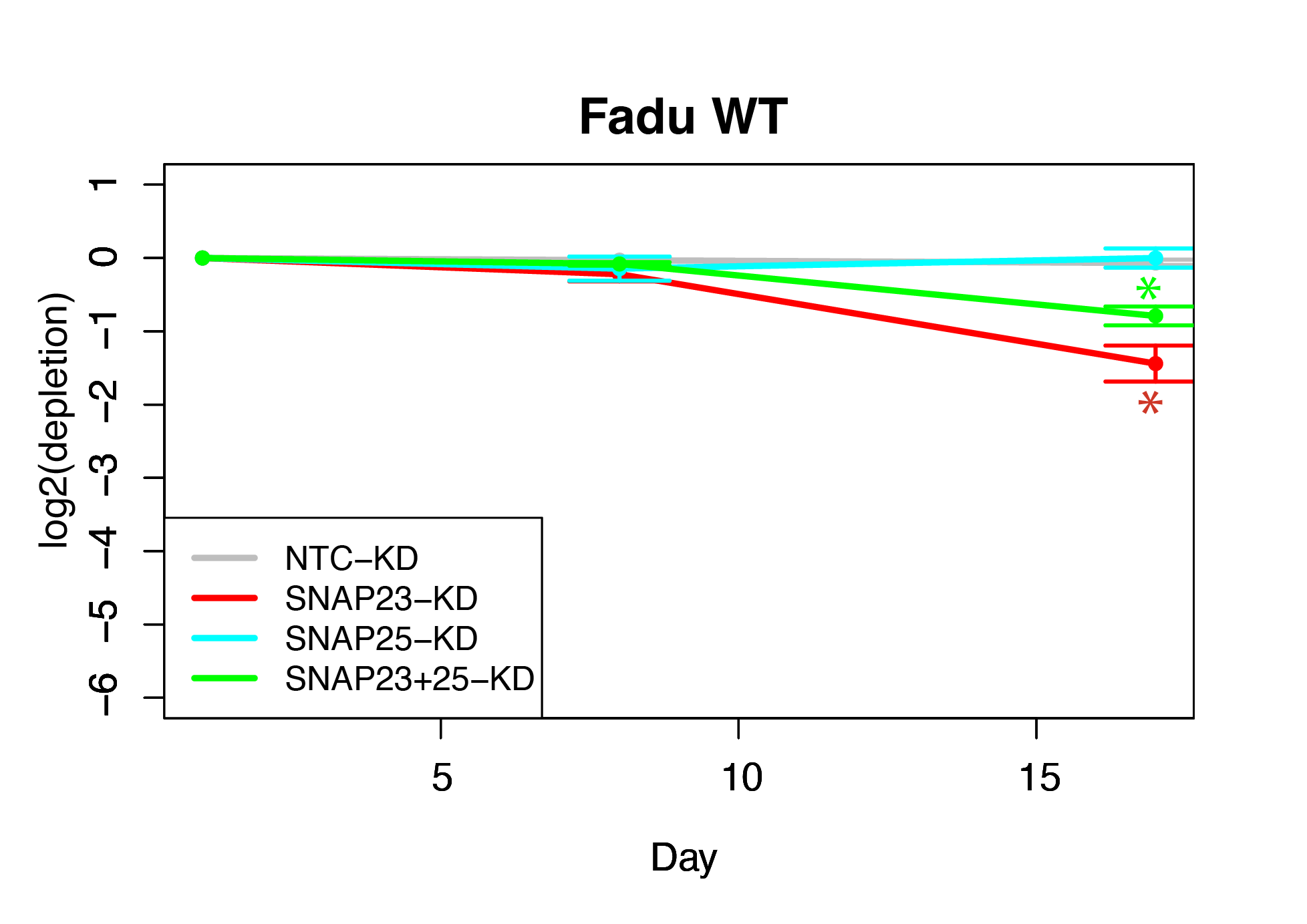

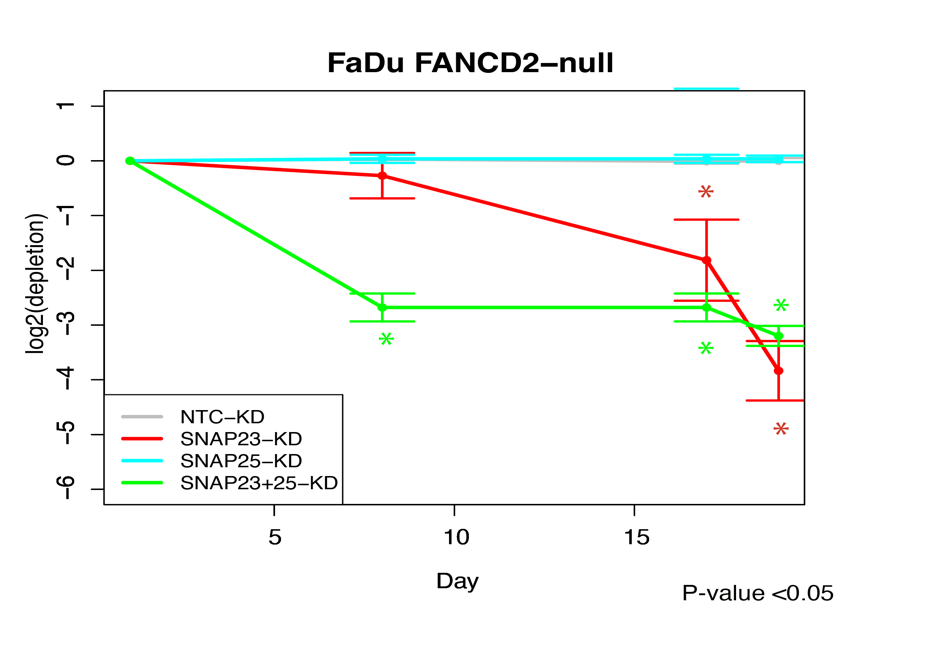

Figure S6) Analysis of various trafficking (25/27 genes) and general autophagy genes the FANCD2-null CRISPRi screening results show no significant effects in depletion score. ATG16L1(gene necessary for autophagy by regulating membrane trafficking^10^) had ~-0.96 depletion score while Rab5A (protein that regulates the movement of vesicles by facilitating internalization and fusion to early endosomes^11^) had ~0.63 depletion score. Plot of depletion score vs -log10(adjusted p-value).

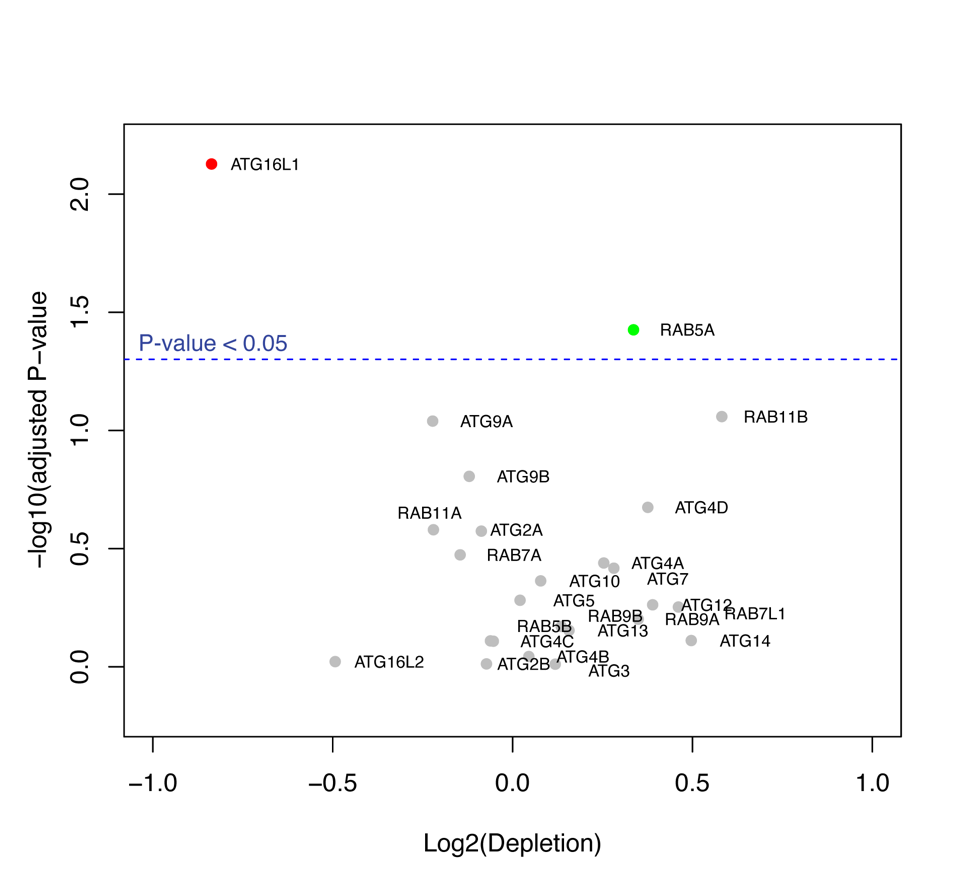

Figure S7) Lysosomal exocytosis was activated using a chemical (ML-SA1) that activates MCOLN1 in FaDu FANCA/D2-null cells. Surface LAMP-1 scars were then quantified using antibodies and flow analysis. Lysosomal exocytosis can be activated with ML-SA1 in FA loss of function mutants and WT. The data for FaDu WT, FANCA-null, and FANCA-null +TG is replotted with ML-SA1controls from Figure 2a.

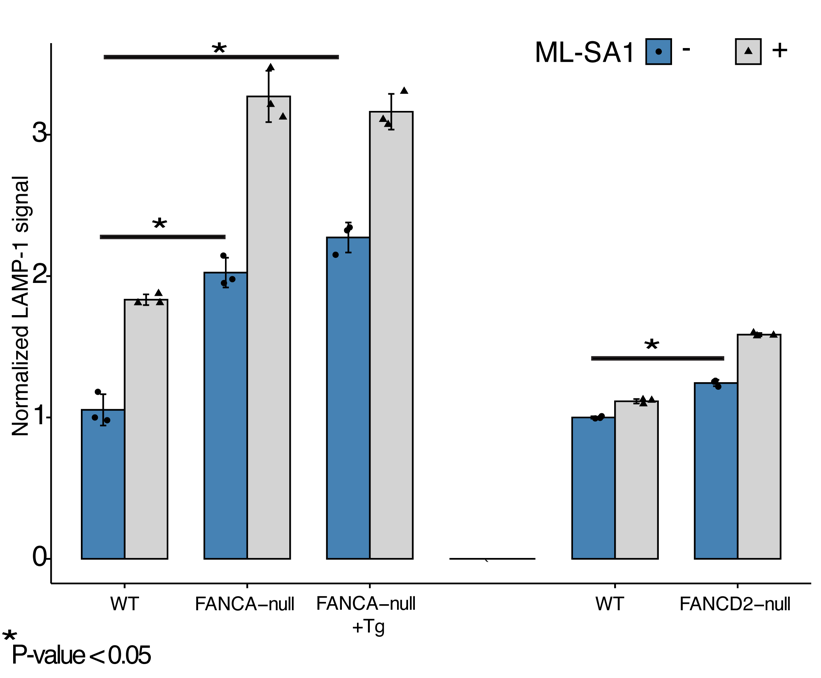

Figure S8) Lysosomal exocytosis was quantified using antibody binding surface LAMP-1 for head and neck cell line, UM-SCC-01. LAMP-1 scars on the plasma membrane were quantified by flow analysis (n = 3 as biological replicates; unpaired t-test was used to determine statistical significance).

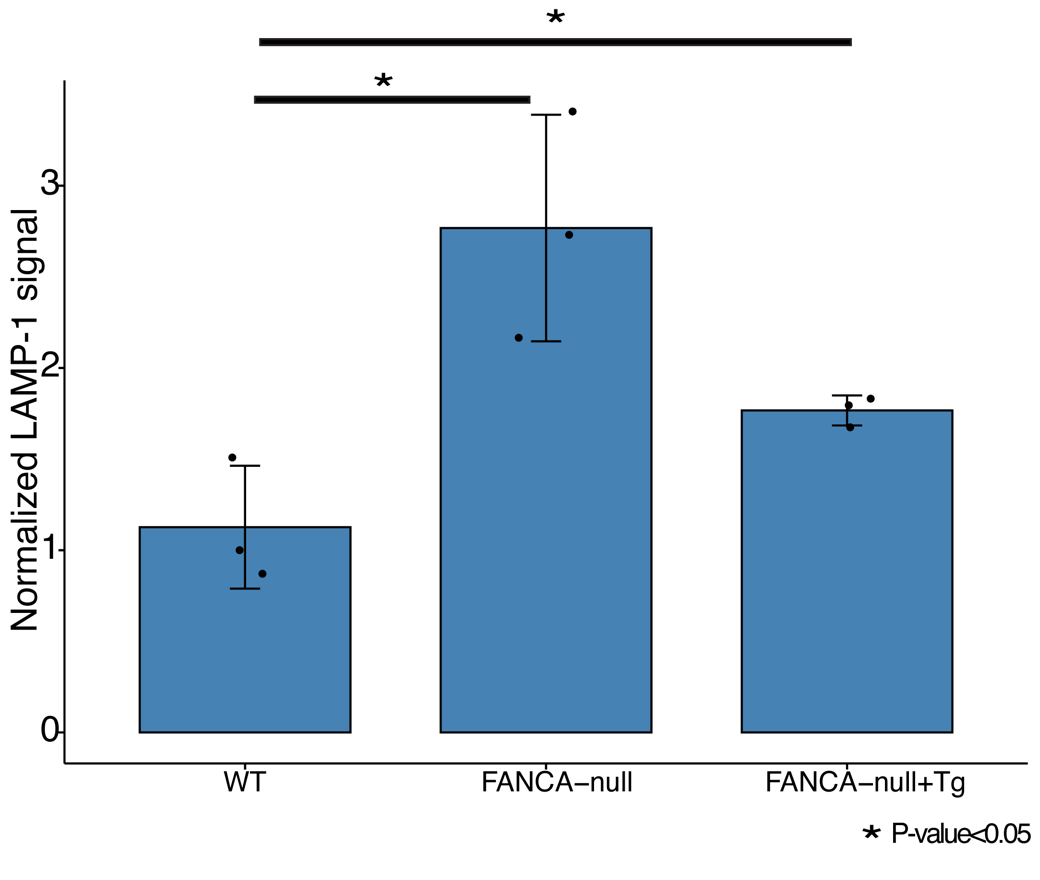

Figure S9) Immunoblot quantification shows the rescued FANCA-null cell lines for FaDu and UM-SCC-01 have higher expressions of FANCA than WT. (n = 3 as biological replicates).
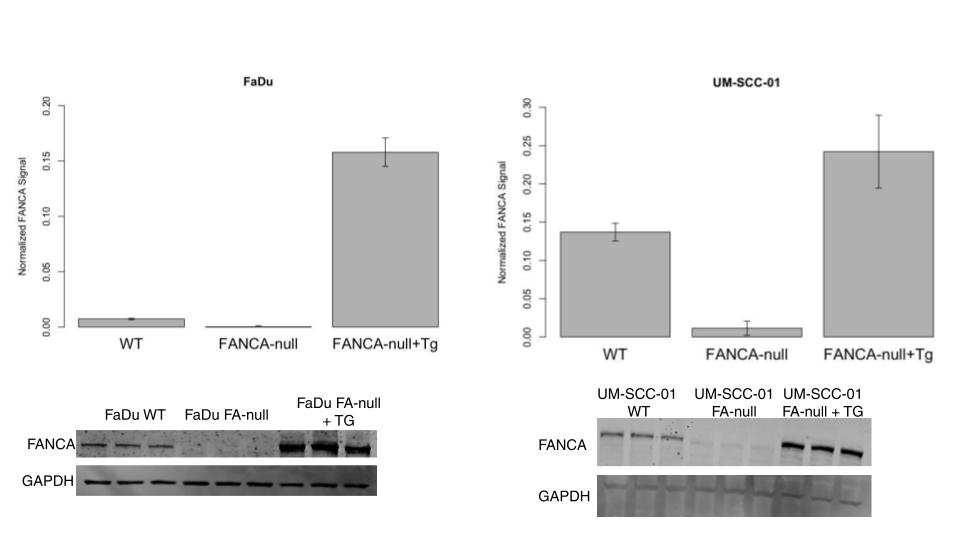

Figure S10) LysoTracker dye retention defects in UM-SCC-01 WT, FANCA-null, and FANCA-null+Tg. LLoMe was used was used as LMP inducing agent and shows that LMP manifests in the UM-SCC-01 cell line as lysosomes enlarging and thus holding more dye. Therefore, the increase in the dye retention and fluorescent geometric mean means more lysosomal damage that may be due to LMP.

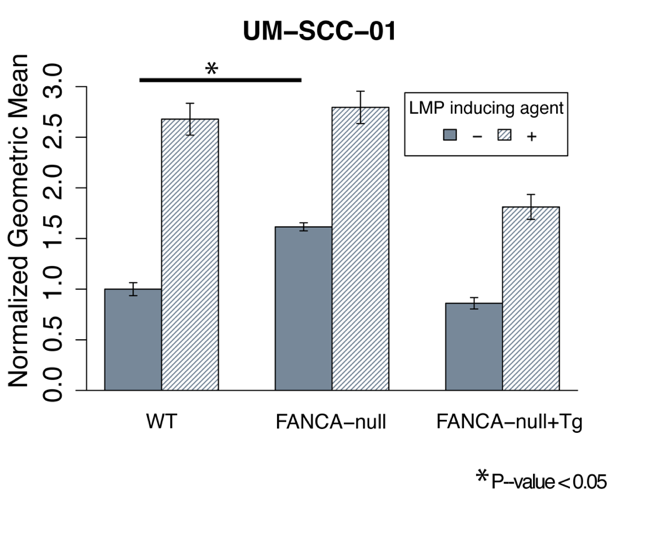

Figure S11) LysoTracker dye was used to quantify LMP and lysosomal health with SNAP23 overexpression (LMP inducing agent LLoMe). (n = 3 as biological replicates; Median ± STD, Unpaired two-tailed t-test was used to determine statistical significance). SNAP23 was overexpressed by integrating SNAP23 via lentiviral integration.

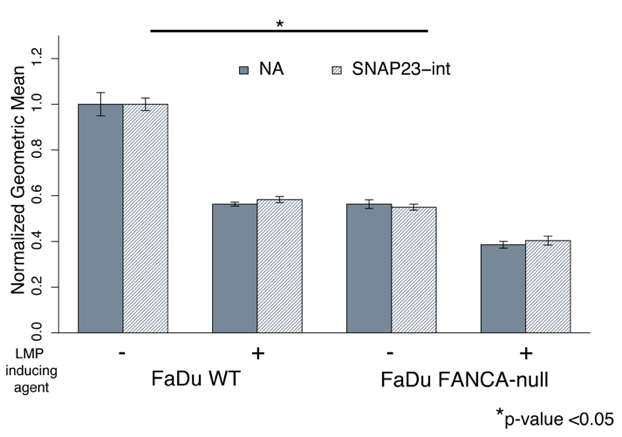

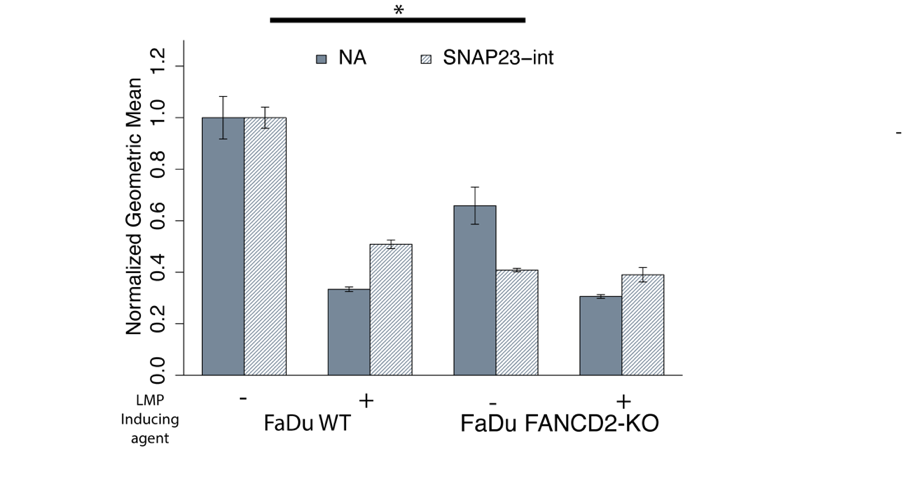

Figure S12) Lysotracker assay was also performed on various cell lines. The LMP inducing agent for the cell lines FaDu, Cal33, and LN18 showed that lysosomal damage manifests in the lower retention of the dye. The loss of FANCA in FaDu and Cal33 and the repression of FANCD2 in LN18 showed decreased ability to retain the LysoTracker dye. The LMP inducing agent in UM-SCC-01 and LN229 showed lysosomal damage manifests in higher dye retention. The loss of FANCA in UM-SCC-01 and the repression of FANCD2 in LN229 showed a trend of increase in LysoTracker dye retention. Lysosomal defects were assessed for control (WT cells or NTC) or FANC gene knockout or knockdown conditions in FaDu, Cal33, UM-SCC-01, LN18, and LN229 as indicated. The loss or repression of FA pathway (FANCA or FANCD2) in the cell lines FaDu, Cal33, LN18, and UM-SCC-01 show statistically significant changes in LysoTracker fluorescence (-1.61, -3, -4.17, +1.62-fold, respectively). LLoMe was used as LMP inducing agent.

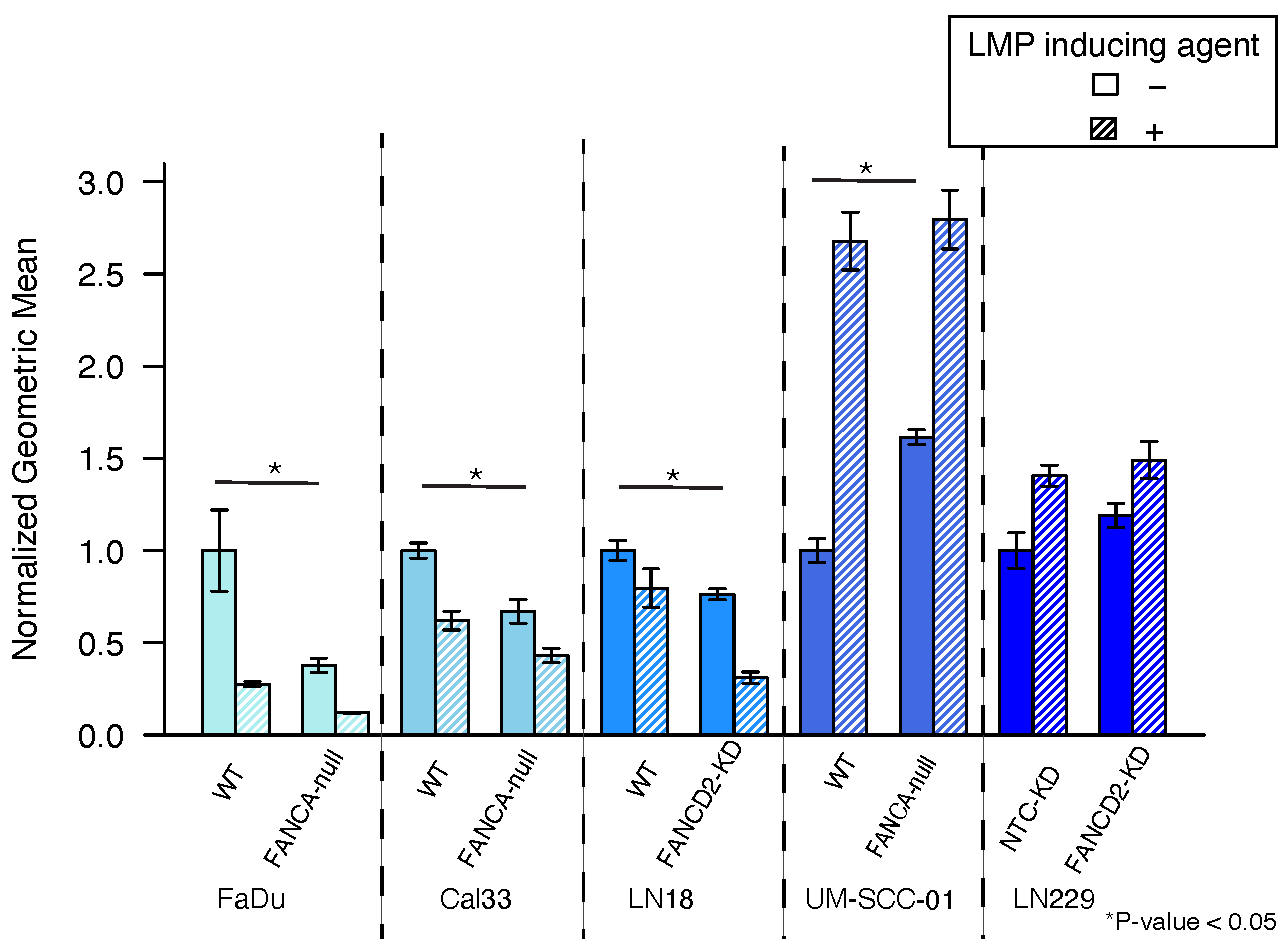

Figure S13) The non-essential genes: FANCA/B/D2/F, were tested in RPE-1 CRISPRi cells. FANCA and FANCD2-KDs were found to have a statistically significant effect on dye retention (~1.67 and 2-fold increase, respectively). LLoMe was used was used as LMP inducing agent.

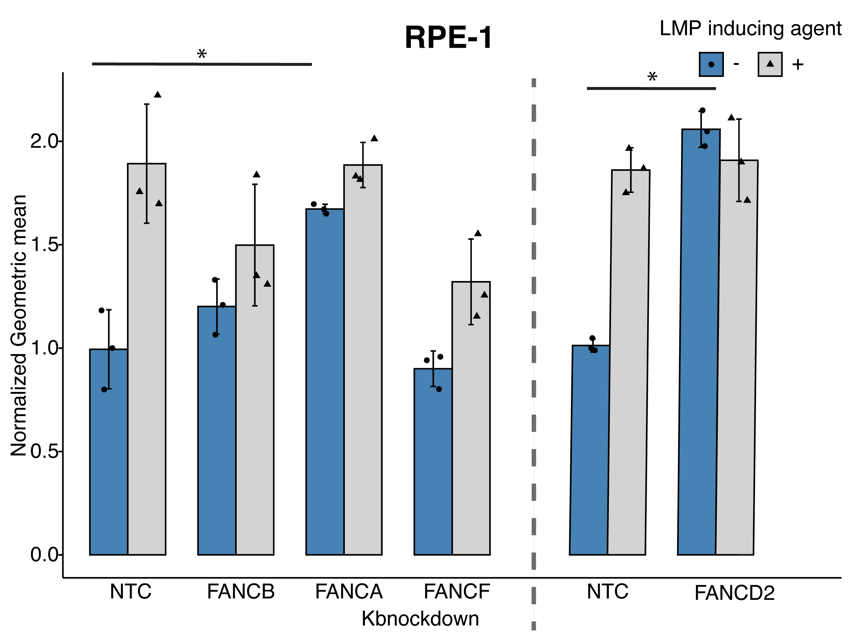

Figure S14) LysoTracker dye was used to quantify LMP and lysosomal health with following DNA repair related candidates (FANCA, XRCC4, FBXO42, ATMIN, BRD8) and negative controls (NTC, HUWE1) in the FaDu CRISPRi cell line. All genes used for knockdown had a depletion score less than -1 or had no effect in the screen. We did not want to assay any candidate that had significant death. The LMP inducing agent used is LLoMe (n = 3 as biological replicates; Median ± STD, Unpaired two-tailed t-test was used to determine statistical significance).

The FANCA knockdown served as a positive control while HUWE1 (an E3 ligase) served as negative control. FANCA-KD showed a decrease in ability to hold the dye, but the phenotype is not as high compared to WT. This difference is likely due to incomplete knockdown. HUWE1-KD, as expected, did not show a difference in the LysoTracker assay. XRCC4 is a DNA ligase that rejoins DNA strands in NHEJ. ATMIN is required for ATM-mediated signaling and recruitment of 53BP1 to DNA damage sites. BRD8 has been shown to be required for DNA repair and preventing DNA damage. FBXO42 is a ubiquitin ligase that is important for mitosis and can cause cellular arrest and spindle dysfunction which can lead to replication stress in cells.

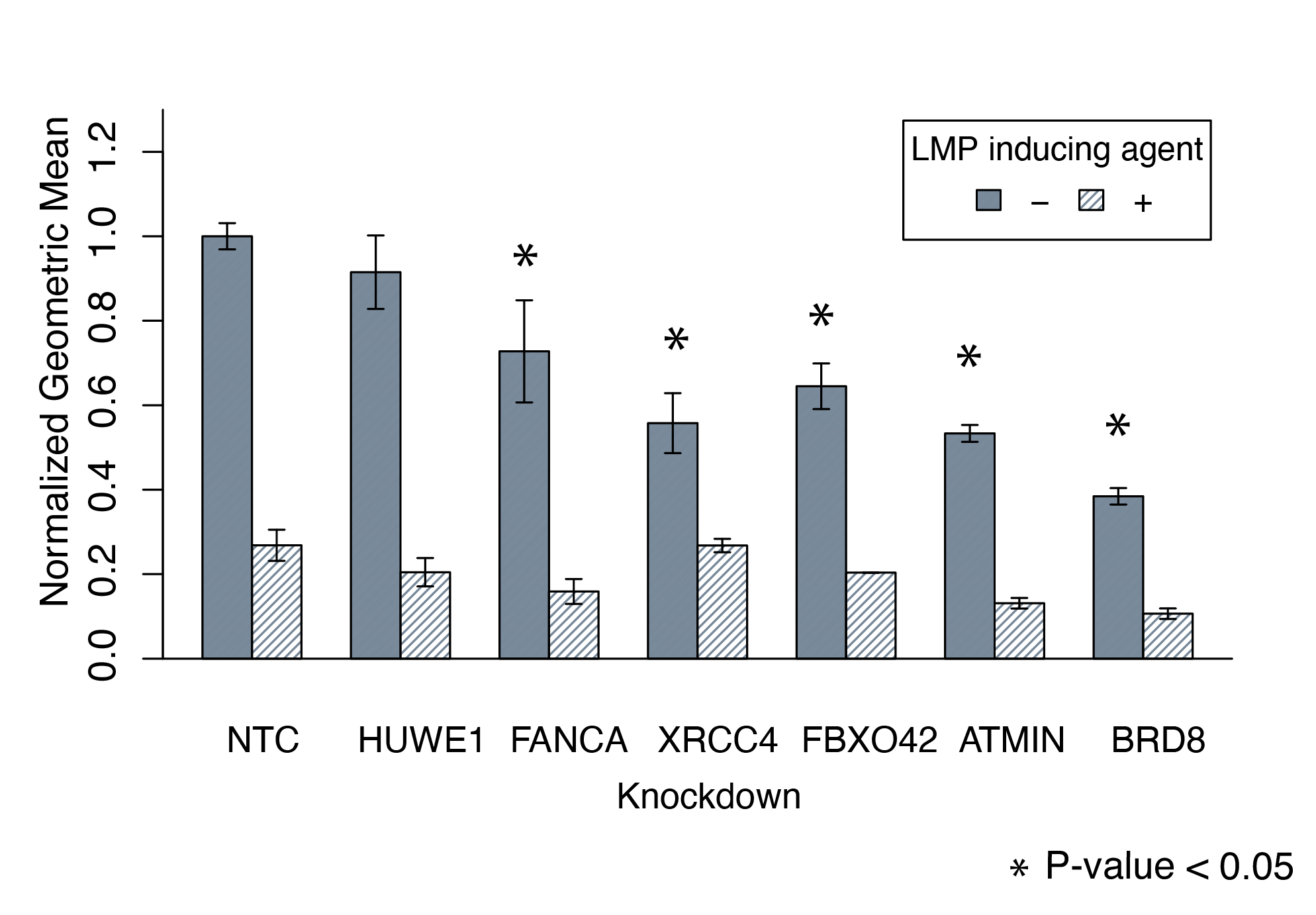

Figure S15) Lysosensor and DQ-BSA Red was used to characterize the lysosomal defects in cells with the loss of function in FANCA or FANCD2.

a) The LysoSensor yellow/blue dye was used to assay pH of cells in FaDu and UM-SCC-01 with and without FANCA or FANCD2. The pH of the lysosomes is more basic in FA mutants in both cell lines.

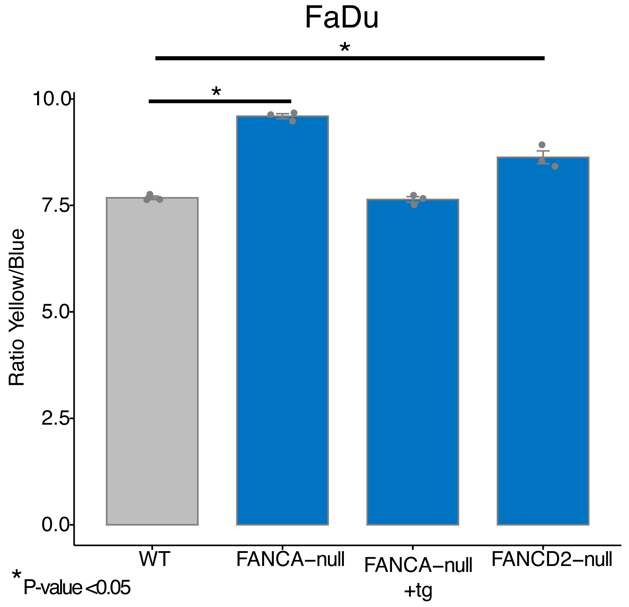

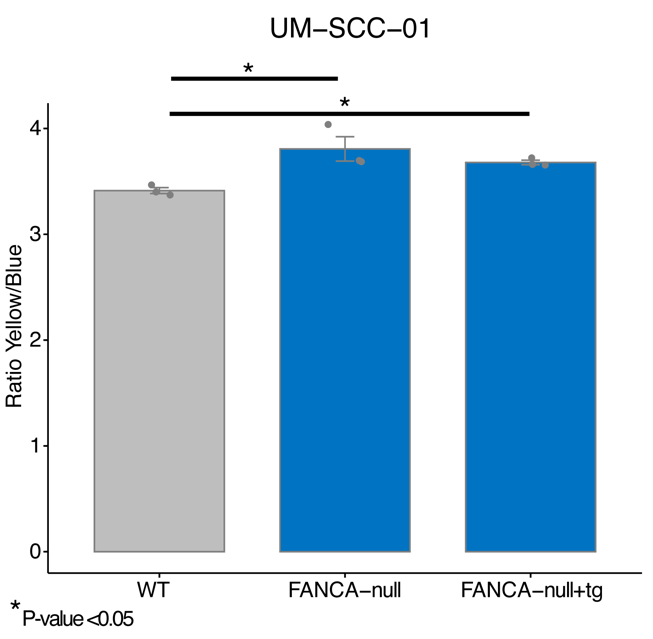

b) DQ-BSA Red is a fluorogenic substrate for proteases that can used to study the delivery of cargo to the lysosomes and monitor autophagy. When DQ-BSA Red is in an acidic environment, proteases cleave the fluorescent peptide. The lysosomal activity of FANCA-null is slightly higher in FaDu and UM-SCC-01 FA loss of function mutants.

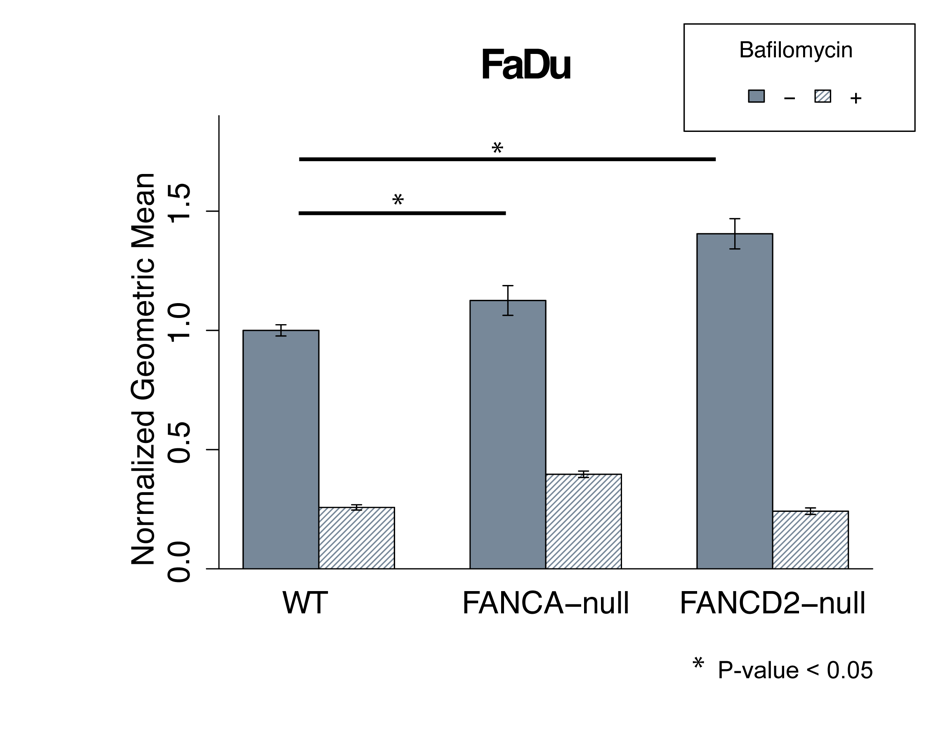

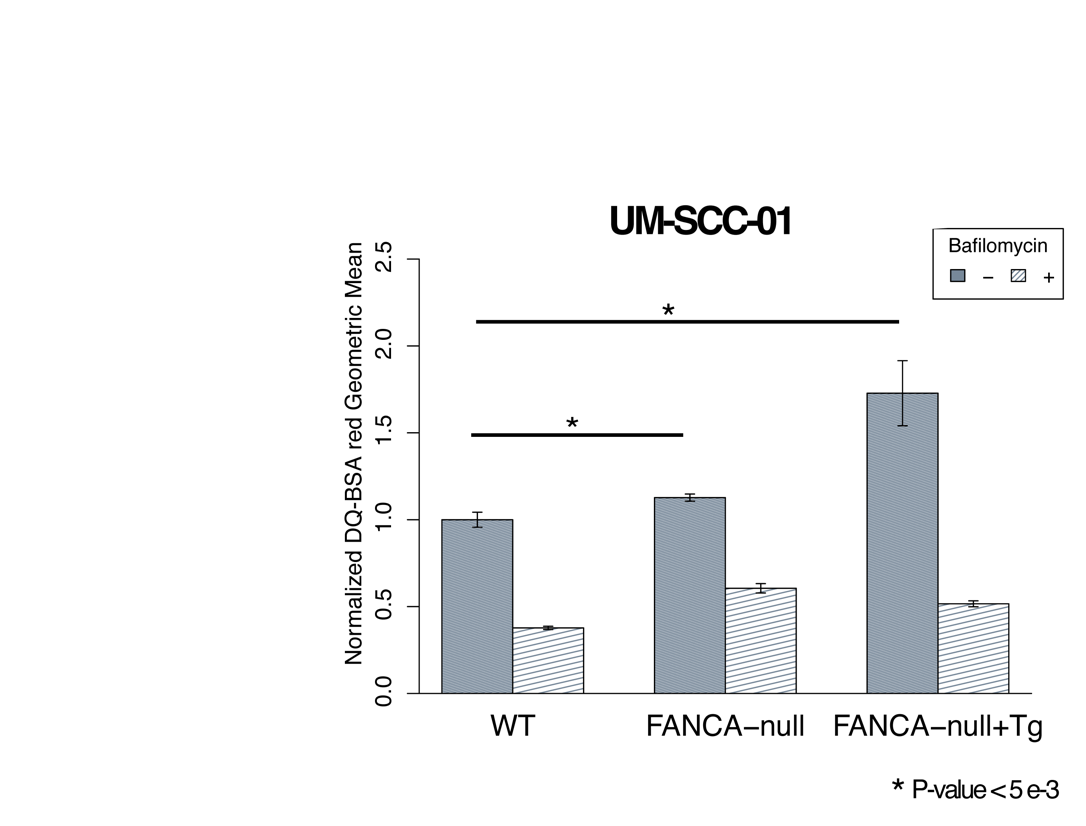
Figure S16) Pulse chase experiments with Dextran-FITC were used to assess LMP in FANCA UM-SCC-01 cells. Dextran-FITC is loaded into lysosomes and chased with fresh media. Cells undergoing LMP will release Dextran-FITC into the cytosol which will result in increased fluorescence as seen in the +LMP inducing agent (LLoMe) conditions. FANCA-null mutants show an increase in cytosolic Dextran-FITC release in UM-SCC-01 cells.

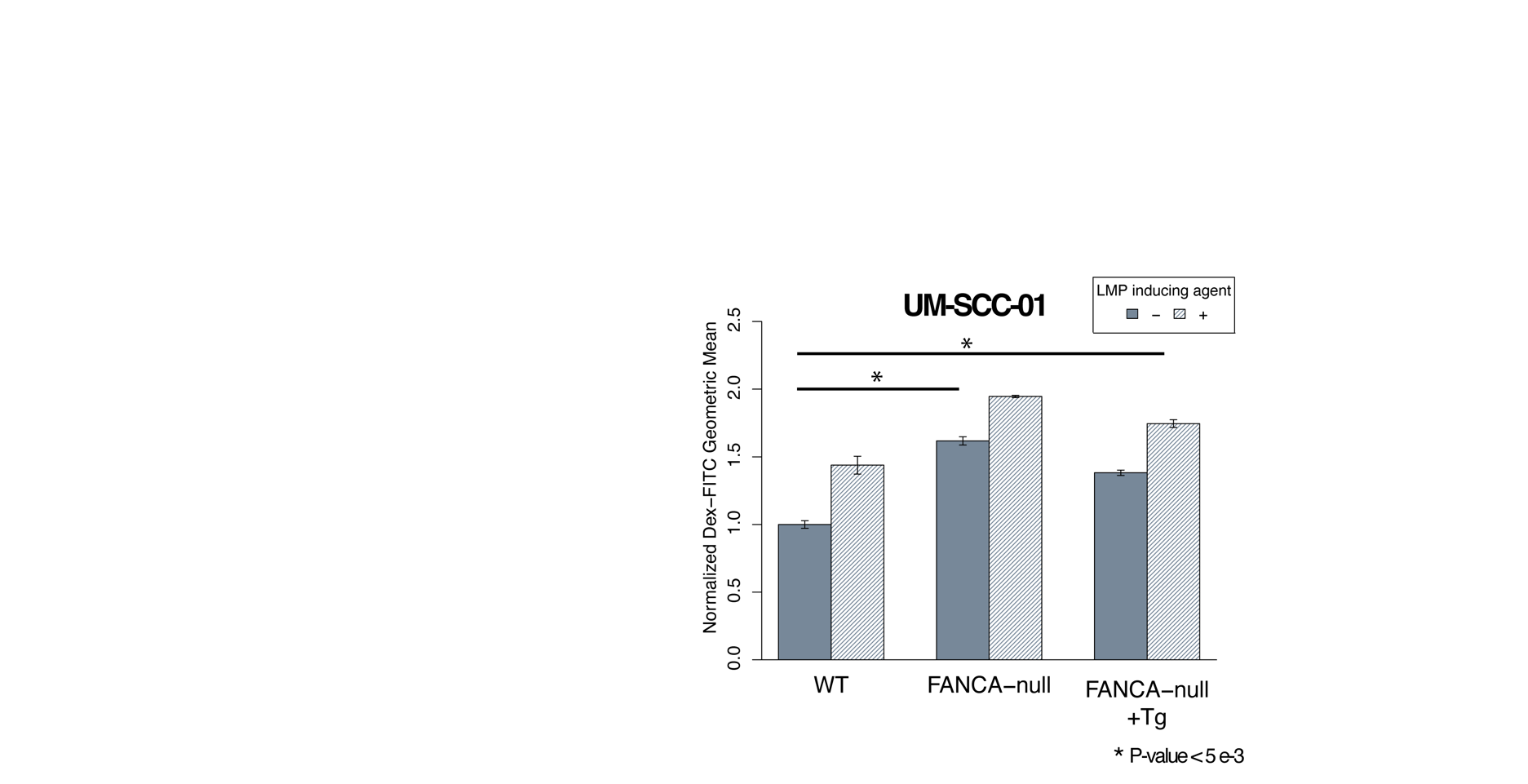

Figure S17) Lyso-IP was performed in FaDu FANCA/D2-null cells and quantified for LGALS3. The overall number of lysosomes in the Fanconi anemia mutants are lower (see Figures 2-3) thus the LGALS3 signal was normalized against LAMP-1 signal. Biological replicates were performed for these experiments and representative blots are shown.

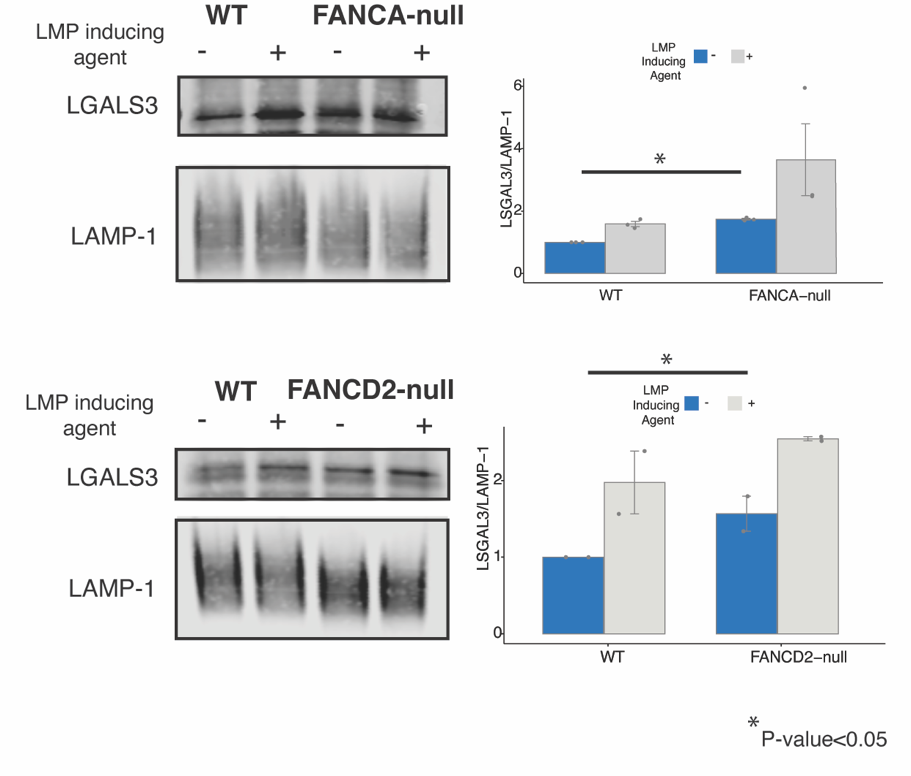

Figure S18) LysoIP was performed to isolate lysosomes from WT, FANCA-null, and FANCD2-null FaDu cells. K48 levels were measured using ELISA (LifeSensor PA480) and normalized to LAMP-1 levels (n = 3 as biological replicates; unpaired two-tailed t-test was used to determine statistical significance).

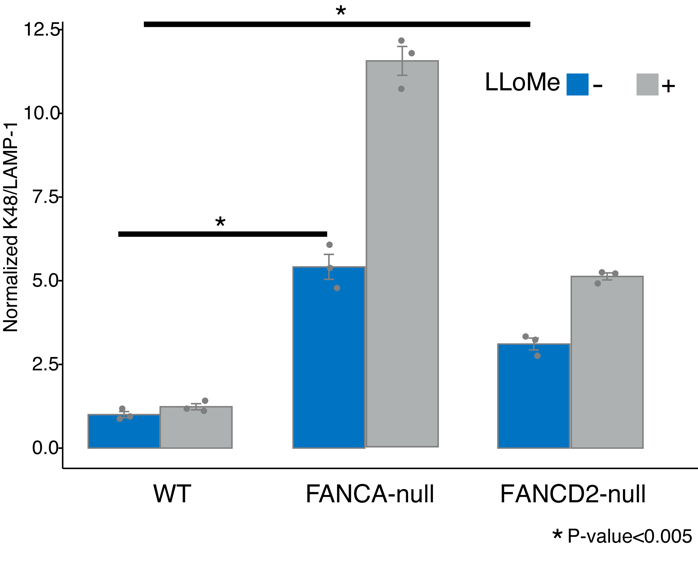

Figure S19) A graph showing clonogenic cell survival as measured by a colony formation assay for WT, FANCA-null or FANCD2-null FaDu, Cal33, and UM-SCC-01 cells were treated with CQ. FaDu FANCA and FANCD2-null showed a 2.6 and 1.78-fold increase in growth inhibition. Cal33 FANCA-null similarly was more sensitive to CQ than WT (~-1.33-fold sensitivity). UM-SCC-01 FANCA-null had a ~-3.85- fold sensitivity that was non-statistically significant. (n = 3 as biological replicates; unpaired two-tailed t-test was used to determine statistical significance).

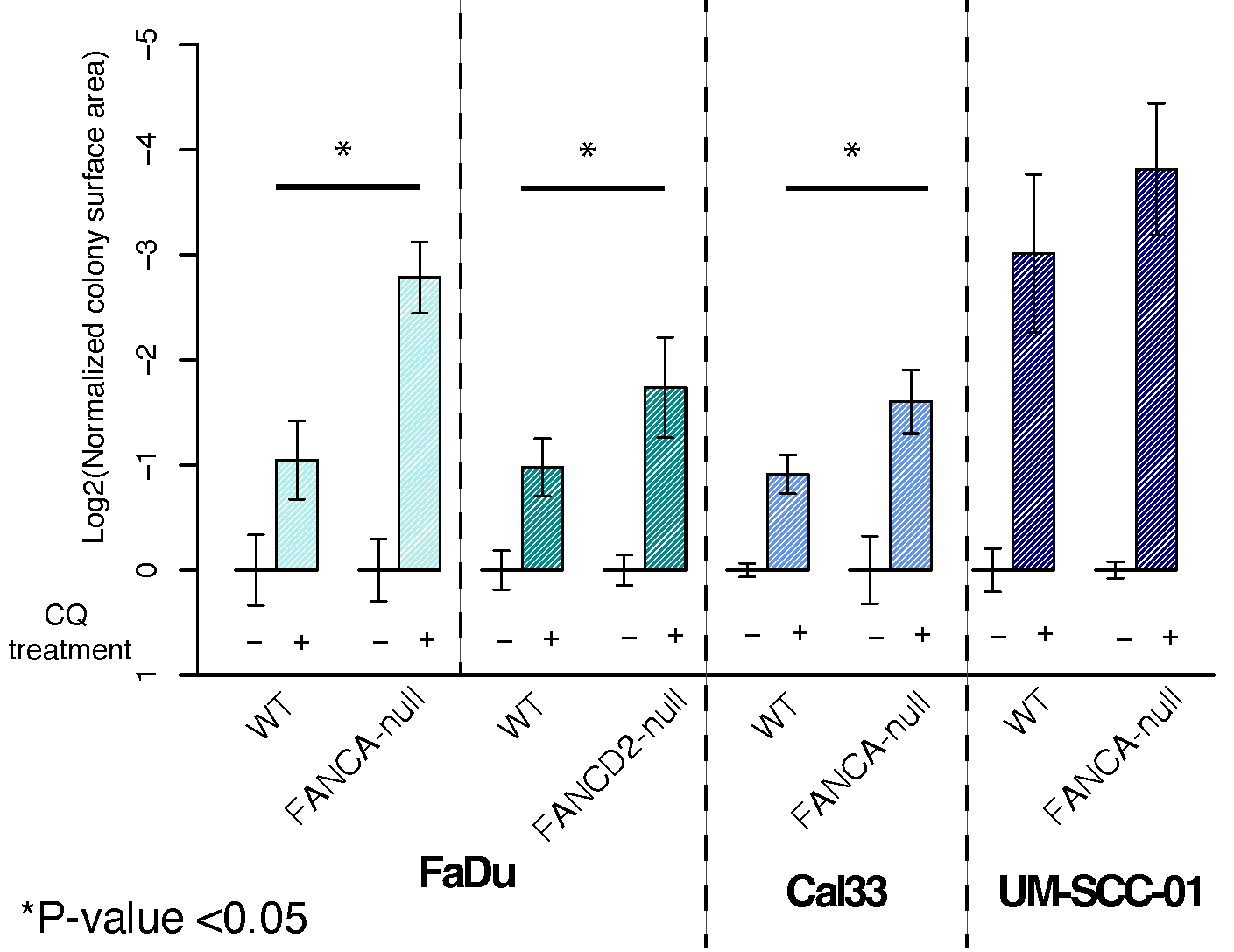

Figure S20) Immunofluorescent data from Figure 3 plotted alongside LMP inducing agent as a control. (Two-sided Mann-Whitney test was used to assess statistical significance)

a) Mean foci intensity data from Figure 3a-c plotted with LMP inducing agent.

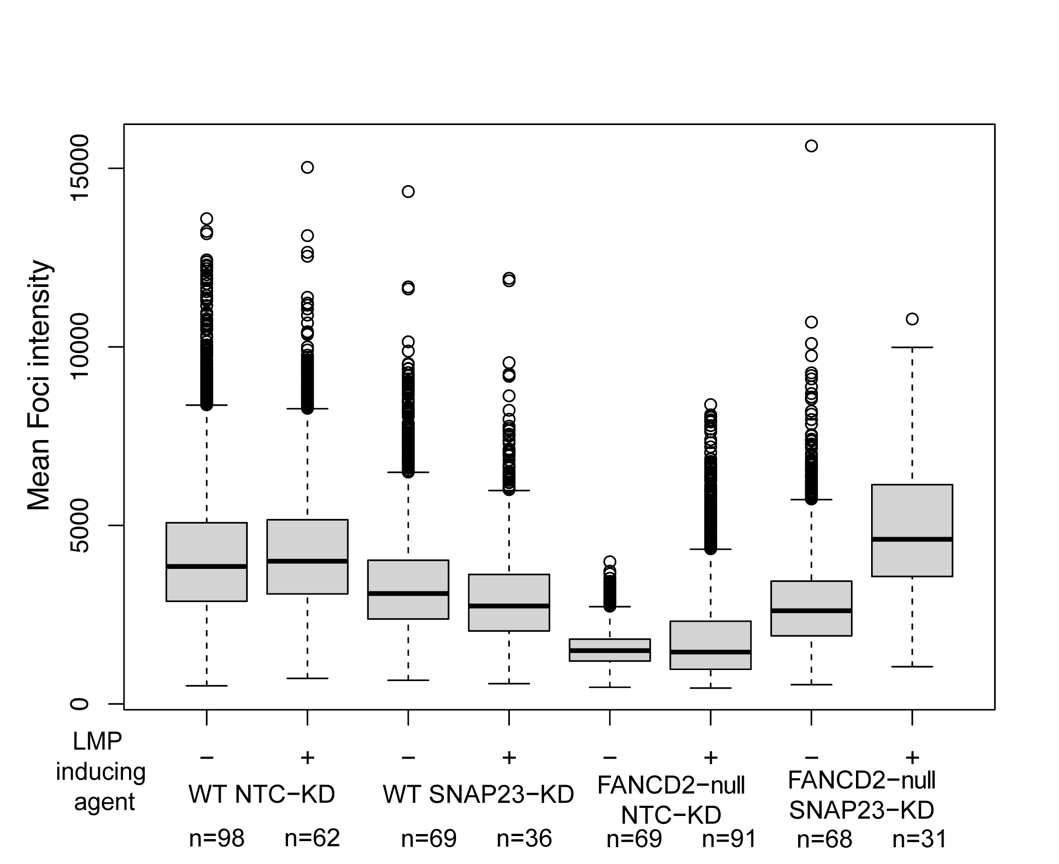

b) Foci count data from Figure 3a-c plotted with LMP inducing agent.

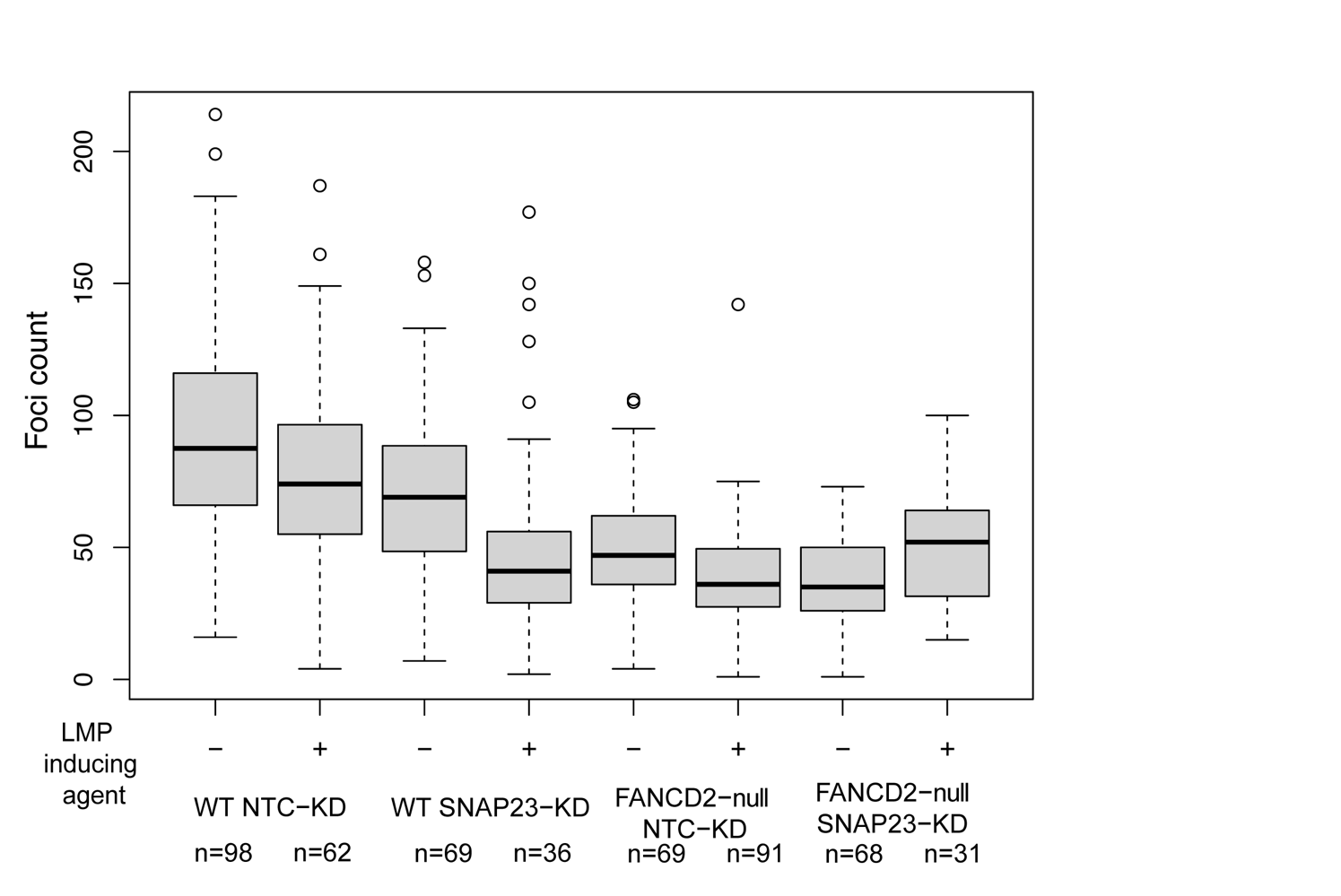
Figure S21) Phospho-4E-BP1/ 4E-BP was used to assess mTOR activation and DNA damage via immunoblotting. 4E-BP1 is a well-known substrate of mTOR. Biological replicates were performed for these experiments and representative blots are shown.

a) FaDu phospho-4E-BP1/4E-BP1 western blot.

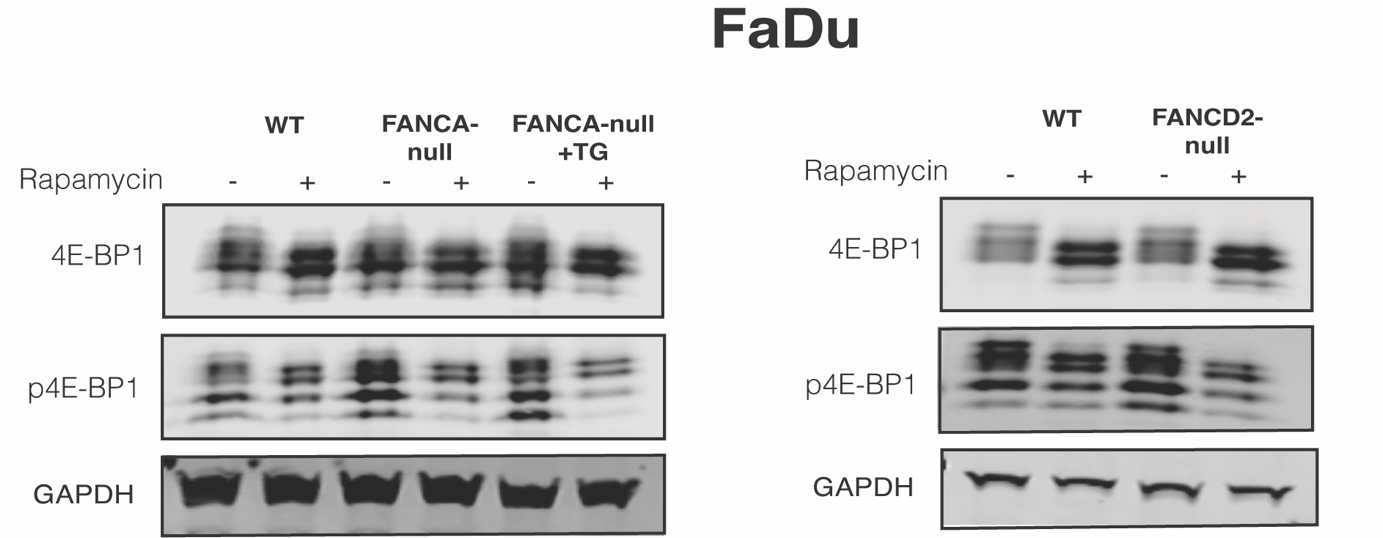

Figure S22) The GFP-LC3-RFP-LC3ΔG assay was also tested in Cal33 WT and FANCA-null backgrounds. The Cal33 FANCA-null had increase in autophagic flux compared to WT with or without SNAP23-KD (p-value > 0.005). (Two-sided Mann-Whitney test was used to assess statistical significance)

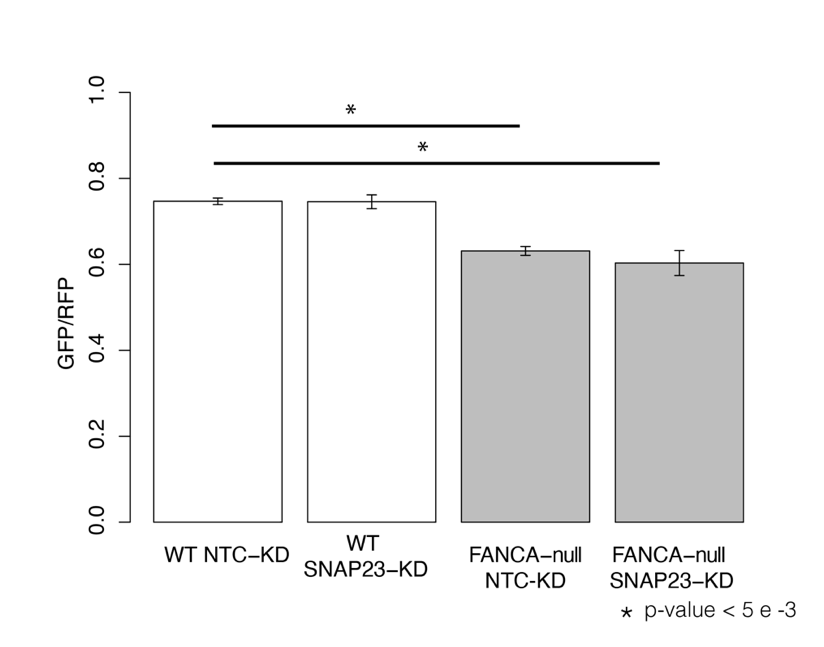

Figure S23) ShinyGO analysis for upregulated genes (p-value > 0.005 & log2(fold difference) > 1.5).

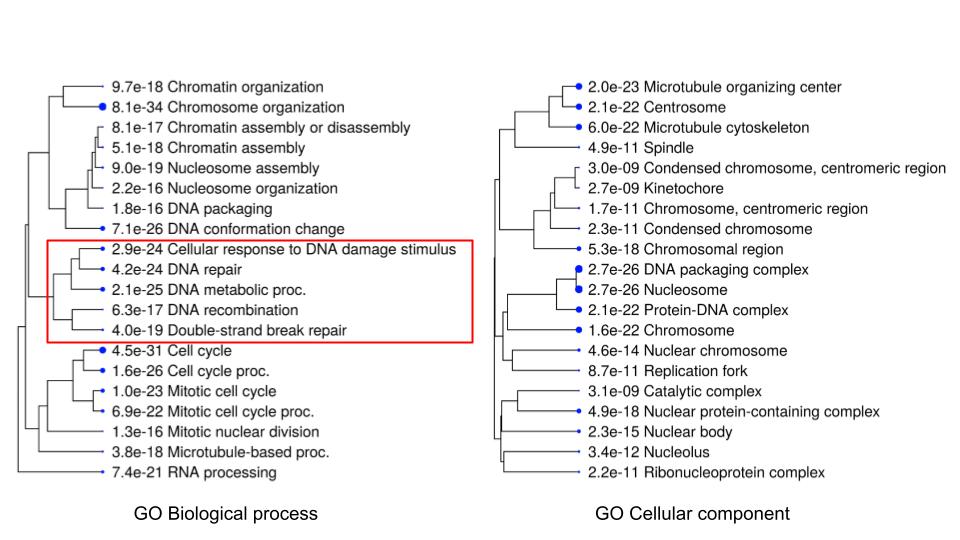

Figure S24) qPCR results for validating the expression level of MCOLN1 in FaDu, Cal33, and UM-SCC-01 head neck cell lines with and without FA knockouts. (n = 3 as biological replicates; Mean ± STD, Unpaired two-tailed t-test was used to determine statistical significance).

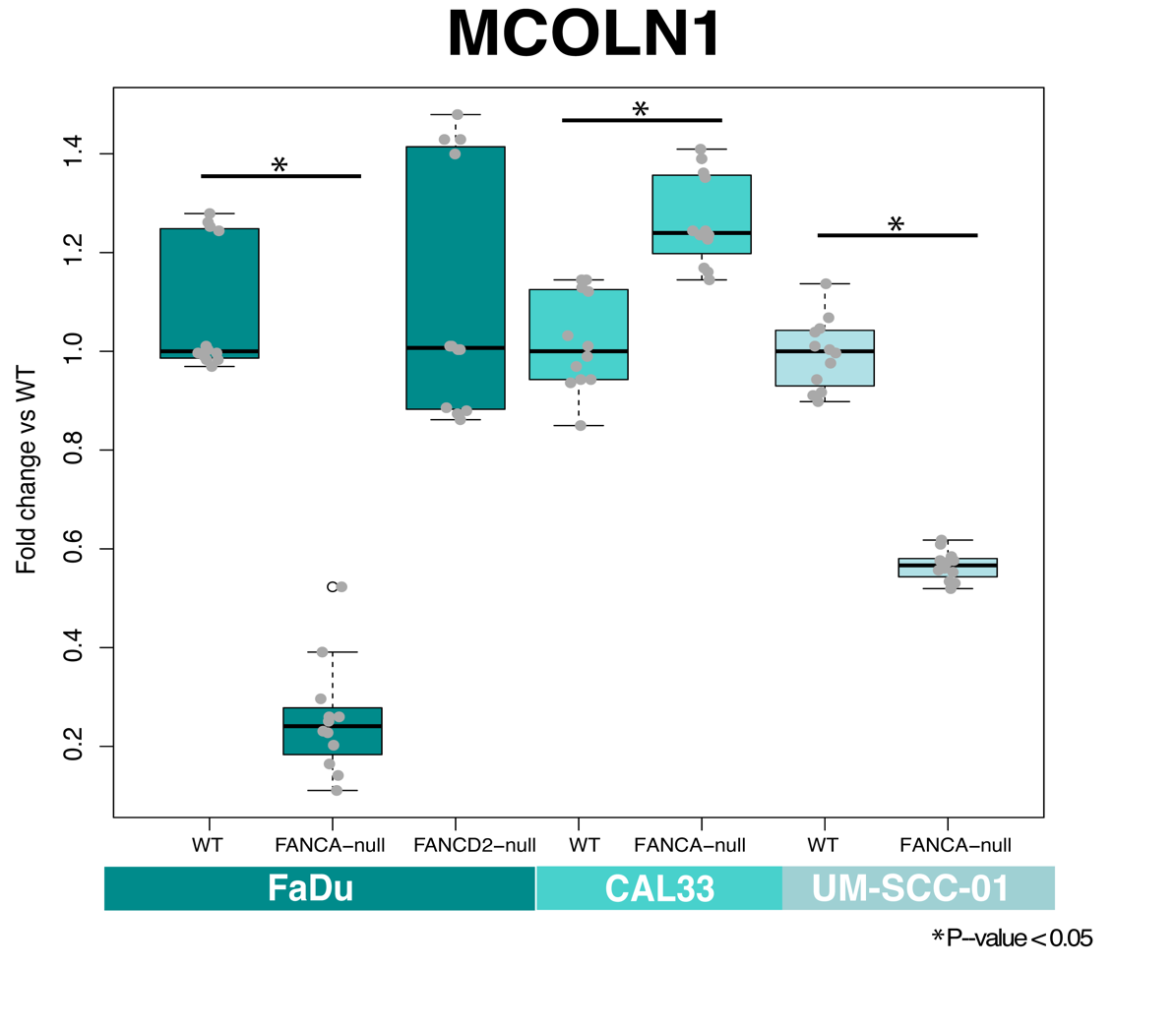

Figure S25) Immunofluorescence was used to quantify TFEB intensity and localization in FaDu and Cal33 cell lines. LMP treatment was used as a positive control to visualize the localization of TFEB when LMP (LLoMe) is induced. (Two-sided Mann-Whitney test was used to assess statistical significance)

a) FaDu FANCA-null cells showed an increase in nuclear/total cytoplasmic integrated TFEB intensity.

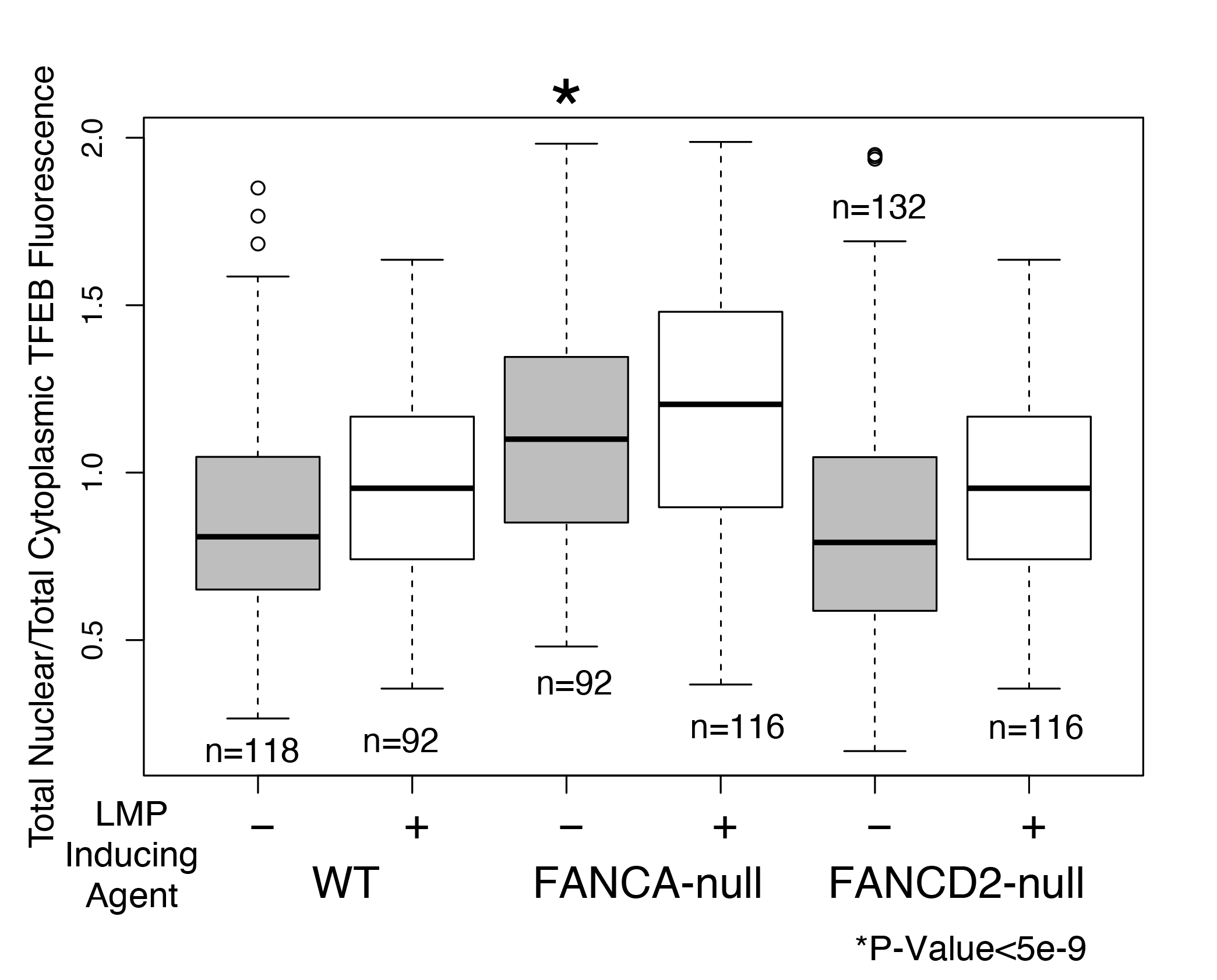

b) The mean TFEB fluorescence intensity confirmed the qPCR results and showed lower integrated TFEB intensity in FANCA/D2-null backgrounds.

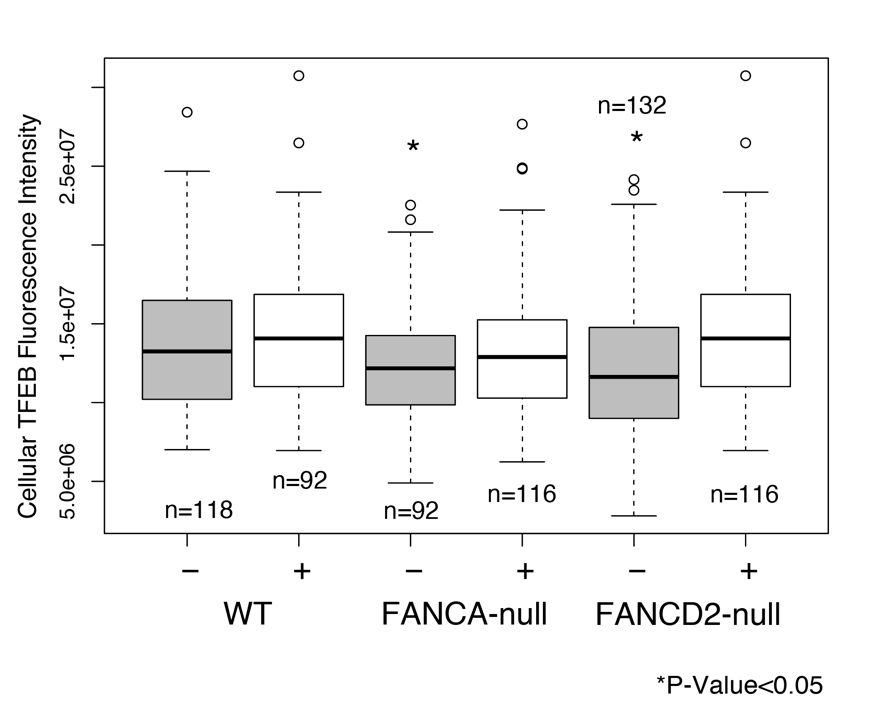

c) Cal33 cell lines showed a lower nuclear/cytoplasmic TFEB mean fluorescence phenotype in FANCA-null. This suggests that Cal33 cells have abnormal TFEB localization that lowers TFEB activation in the FANCA-null background.

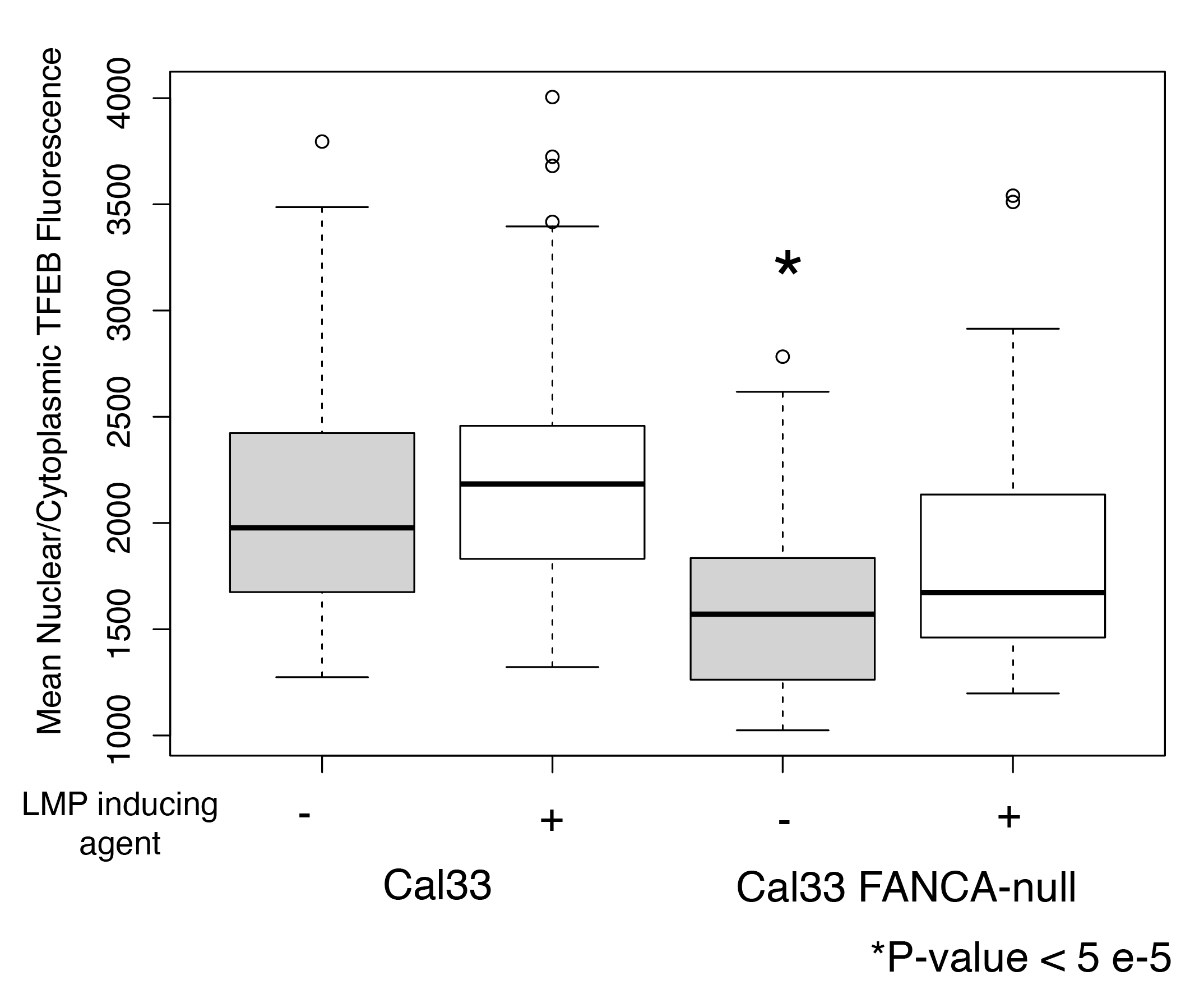

d) Cal33 mean TFEB fluorescence is lower in FANCA-null background which confirms qPCR results.

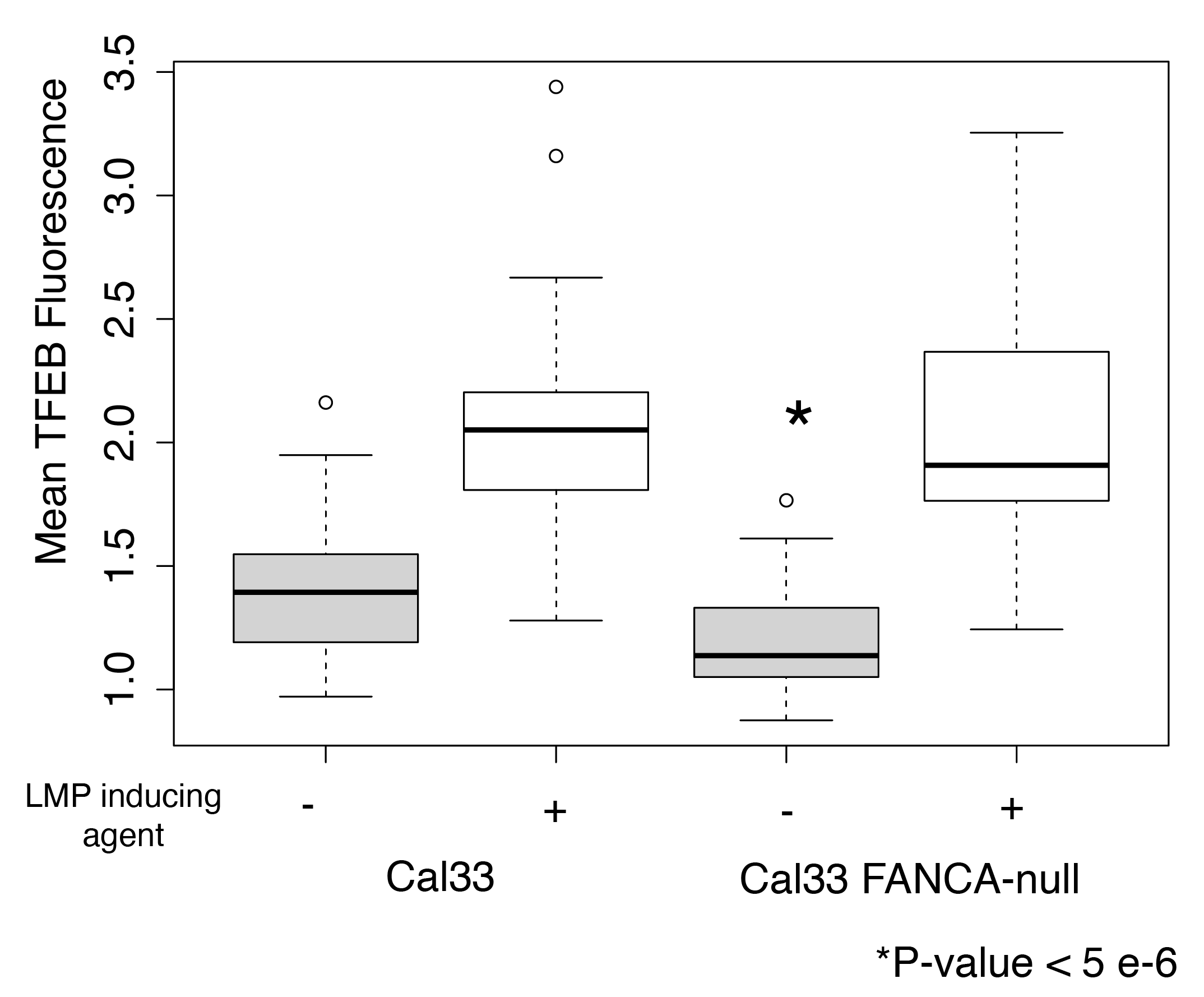

e) LysoTracker dye was used to quantify LMP and lysosomal health with transient TFEB overexpression (LMP inducing agent LLoMe) (n = 3 as biological replicates; Median ± STD, Unpaired two-tailed t-test was used to determine statistical significance).

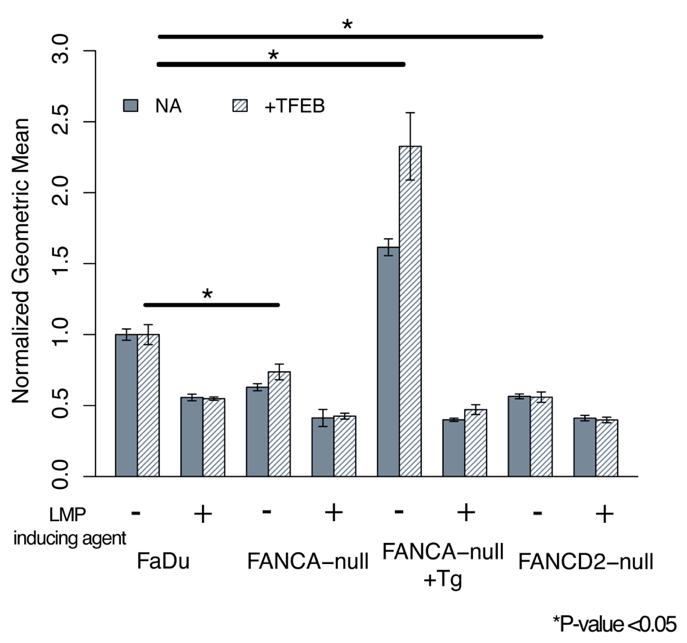

Figure S26) Basal DNA damage was quantified using immunoblotting of H2AX/γH2AX. Biological replicates were performed for these experiments and representative blots are shown. (n = 3 as biological replicates; Median ± STD, Unpaired two-tailed t-test was used to determine statistical significance).

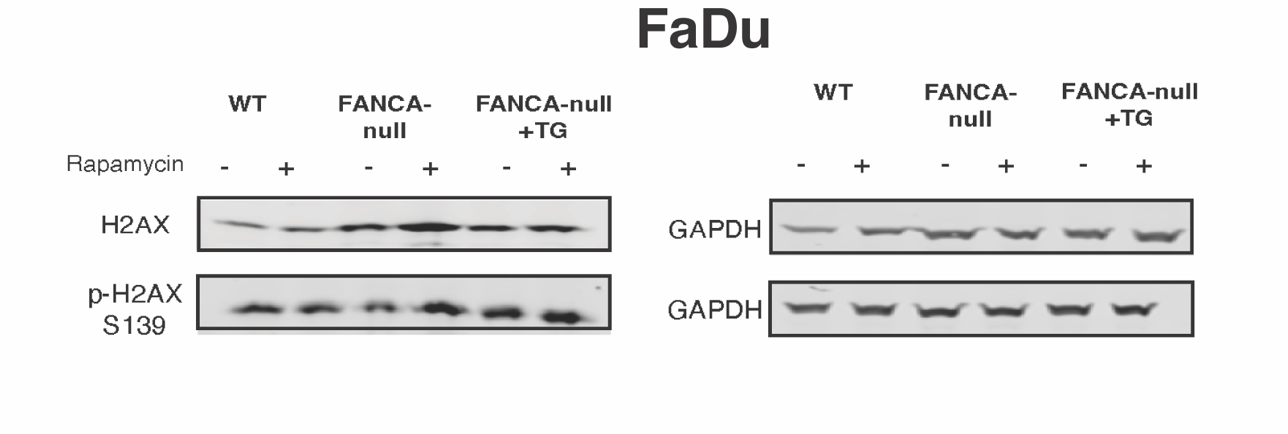

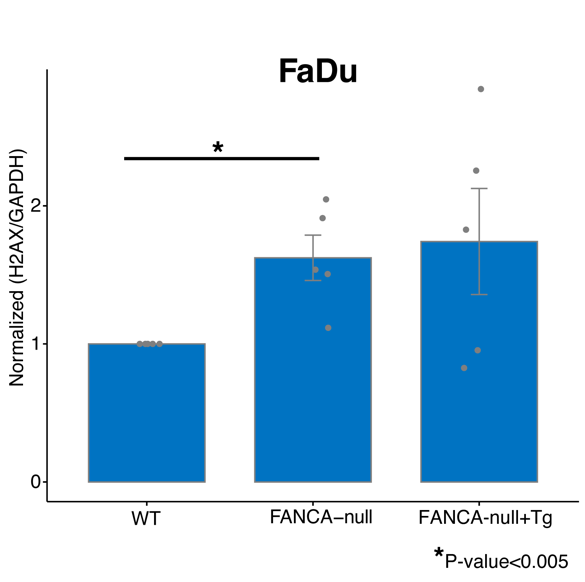

Figure S27) Loss of FANCA (FaDu, UM-SCC-01, Cal33) and FANCD2 (FaDu) cell lines cause an increase in ROS. Grey bars show basal ROS while white bars show 15 min peroxide treated control. (n = 3 as biological replicates; Median ± STD, Unpaired two-tailed t-test was used to determine statistical significance)

a) FaDu cell lines with FANCA/D2-null ROS levels measured using CellROX.

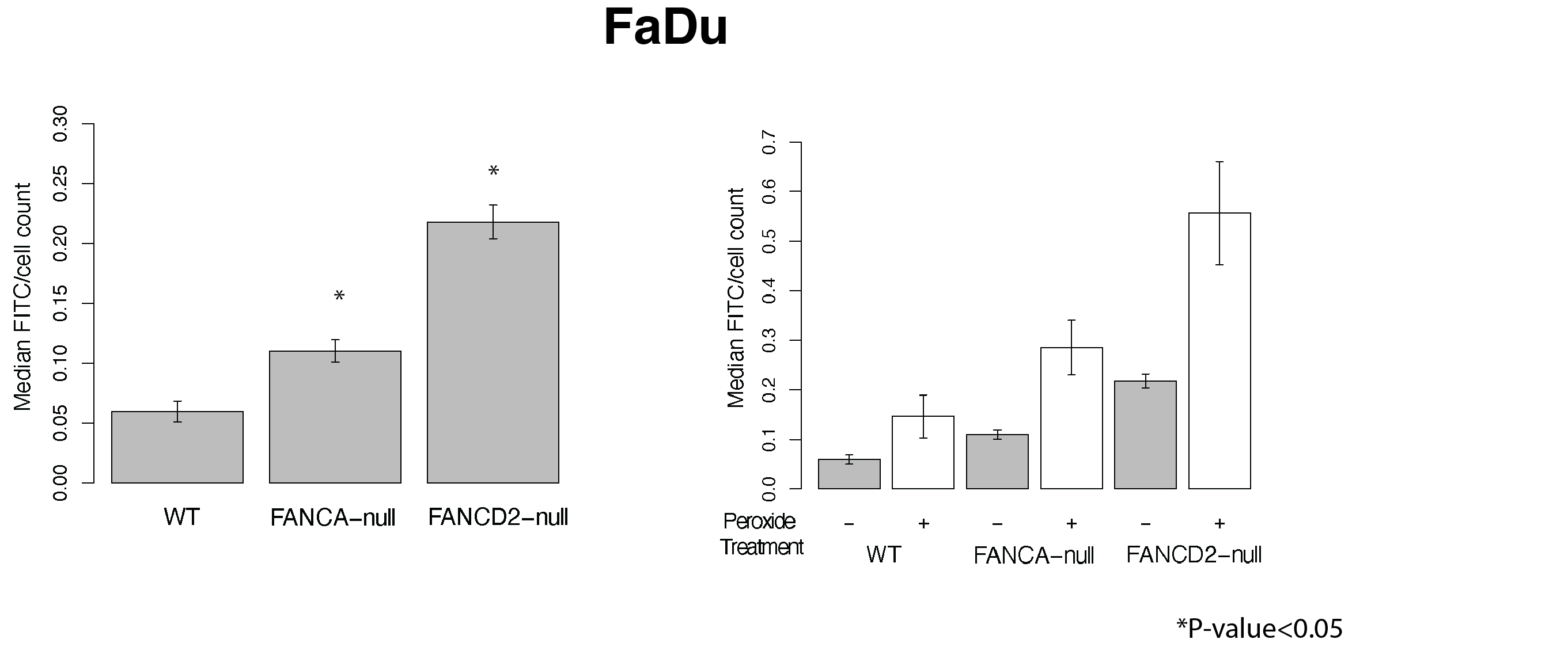

b) UM-SCC-01 cell lines with FANCA-null ROS levels measured using CellROX.

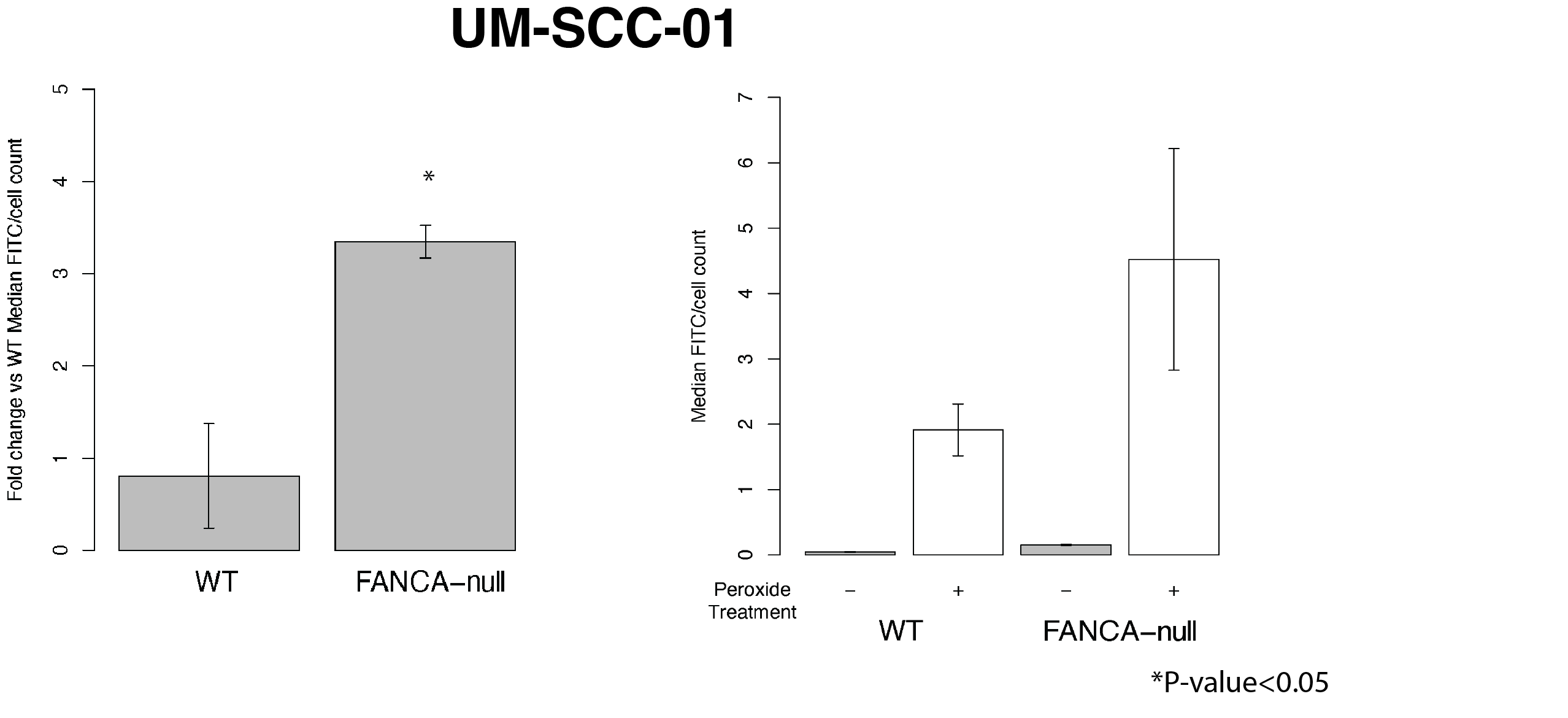

c) Cal33 cell lines with FANCA-null ROS levels measured using CellROX

.
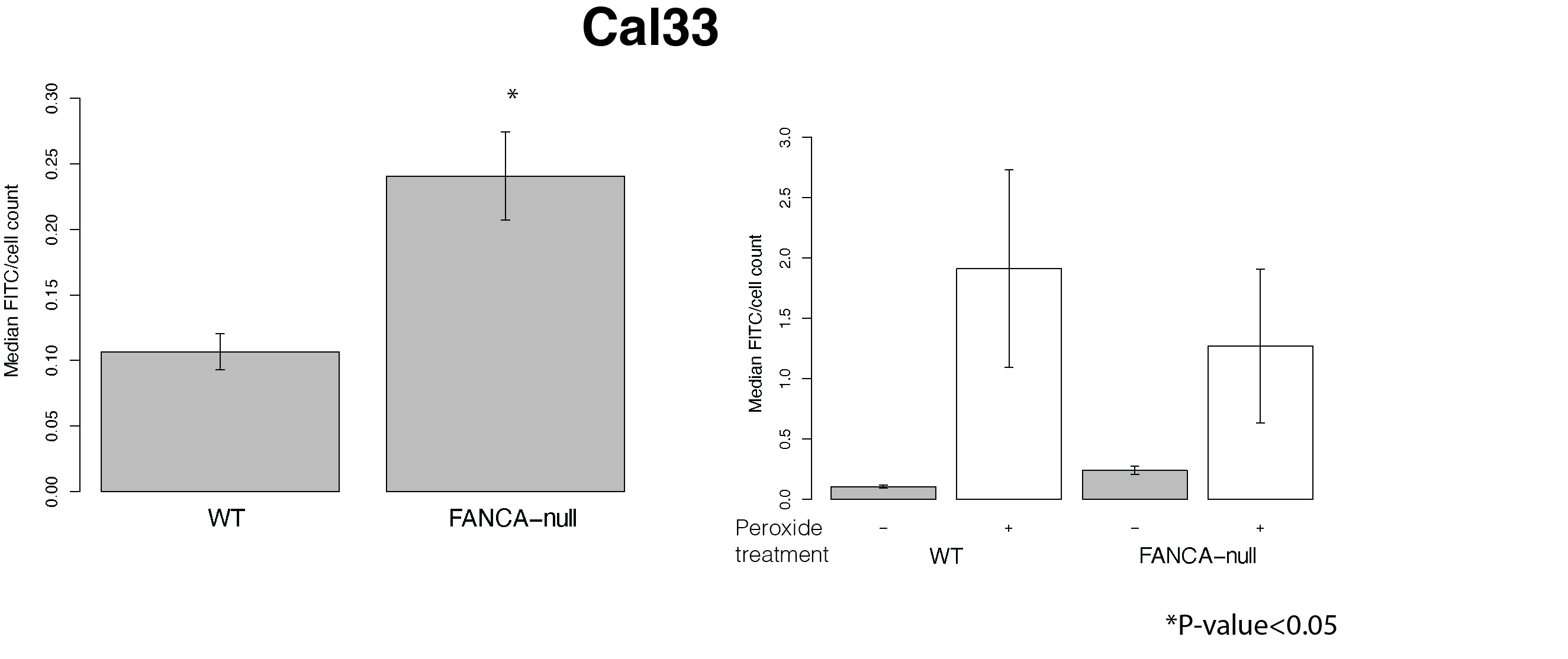

Figure S28) Loss of FANCA (Cal33 and UM-SCC-01) and FANCD2 (FaDu) cell lines cause an increase in mtROS. Colored bars represent basal mtROS while hatched bars show mitoPQ treated positive control. Although the mtROS is not as drastic as overall ROS, a deficiency in the FA pathway shows a trend of increase in mtROS. (n = 3 as biological replicates; Median ± STD, Unpaired two-tailed t-test was used to determine statistical significance)

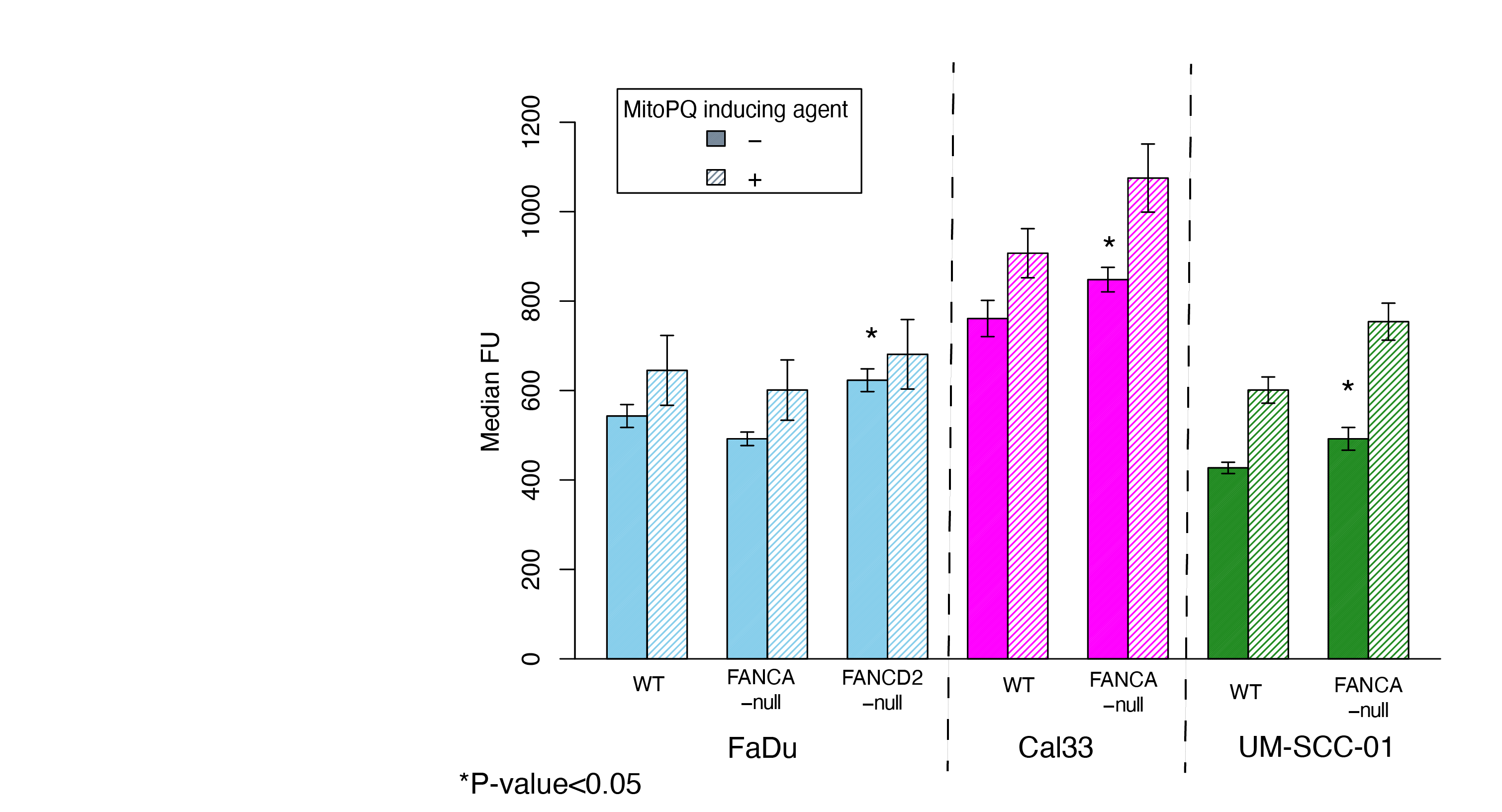
